# Female sex is a risk factor for exacerbated lipid peroxidation and disease in murine retinitis pigmentosa

**DOI:** 10.1101/2025.10.07.680051

**Authors:** Katri Vainionpää, Anna Kalatanova, Umair Seemab, Ahmed B. Montaser, Henri Leinonen

## Abstract

Oxidative stress is an important aspect in retinal degenerations that could be targeted in various forms of currently untreatable diseases. It is generally believed that males are more predisposed to oxidative stress than females due to their higher metabolic activity and/or lower antioxidant capacity. However, studies using mouse disease models have demonstrated that photoreceptor degeneration progresses faster in females. Sex hormones likely play a role, but the cellular mechanism behind the sex difference is unclear. In the current study, we confirmed that the accelerated disease phenotype in female rd10 and P23H retinitis pigmentosa mice coincides with sexual maturity, and further, we found that it co-occurs with increased retinal lipid peroxidation. Instead, protein oxidation and inflammatory marker levels were similar between the sexes. Retinal lipid profiling revealed higher levels of polyunsaturated fatty acid (PUFA)-containing lipids in healthy 2-month-old female mice compared to males, whereas before puberty the sex difference in retinal PUFAs was absent. Analysis of open bulk retina transcriptomic data from middle-aged humans found supplemental evidence of sex-related differences in retinal energy metabolism pathways. Besides mechanistic study directed to reveal the reasons for differential lipid peroxidation between sexes, more research needs to be directed to study sex differences in retinal metabolism and lipid composition across animal species. The current results highlight the need to consider the impact of sex differences when undertaking preclinical experiments with RP models.

**Significance statement:** This article suggests female sex as a significant risk factor for progressive photoreceptor degenerative disease, based on experiments in two widely used retinitis pigmentosa mouse models. The disease phenotype in females accelerates markedly after sexual maturity, especially in mice carrying the autosomal dominant P23H rhodopsin mutation. This acceleration is associated with intensified retinal lipid peroxidation. Given the retina’s high energy demand, continuous photoreceptor cilia renewal, and constant light exposure that generates reactive oxygen species, the susceptibility of photoreceptor membranes to oxidative damage is substantial. Our findings suggest that retinal metabolism may differ between sexes after puberty, potentially influenced by sex hormones, which could contribute to the increased vulnerability of females to photoreceptor degenerative diseases.

## Introduction

The retina is the most energy-demanding tissue in the human body, and particularly its photoreceptors consume large amounts of oxygen [1,2]. The combined impacts of several factors such as: 1) high energy consumption, 2) photo-oxidation of molecules due to light irradiation, 3) the abundance of polyunsaturated fatty acids (PUFAs) that are vulnerable to peroxidation, and 4) the daily shedding of photoreceptor ciliary segments involving the breakdown of lipid-rich materials, render the retina particularly susceptible to suffer from oxidative stress. This vulnerability is further intensified in degenerative photoreceptor diseases. When the highly oxygen-consuming rods die in diseases such as retinitis pigmentosa (RP; 95 and 97 % of the photoreceptors in humans and mice are rods, respectively) [3,4] oxygen demand in the outer retina decreases, but its transport continues at basal levels. This is due to the absence of autoregulation in the choroidal vessels that supply oxygen to the outer retina [5,6]. The resulting hyperoxic environment forms the basis for excessive production of reactive oxygen species (ROS); these radicals participate in secondary degeneration of cones and other collateral damage in the retina [1,2,5–8]. The predisposition to oxidative stress is generally considered to be greater in males and has been speculated to account for their higher incidence of cardiovascular diseases and shorter life expectancy [9–12]. This phenomenon is believed to stem from differences between the sexes in terms of metabolism and antioxidative capacity.

Virtually all common eye diseases manifest a sex-dependent epidemiology [13]. Males suffer from blindness and vision impairment more commonly due to glaucoma and corneal opacity, whereas the vision of females is affected more by diabetic retinopathy and cataract [14]. The lower antioxidant capacity correlates with the lower blood flow to the area of the optic nerve head, which is a marker for the severity of glaucoma, specifically in male patients with normal-tension [15]. It is debatable whether the longer lifespan of females alone explains their higher prevalence of age-related macular degeneration (AMD) [14,16,17]. For example, while early artificially-induced menopause increases the risk of developing this disease [18], hormone replacement therapy is claimed to exert beneficial effects [19]. With respect to inherited retinal degenerations (IRDs), a meta-analysis of *ABCA4*-mutation-affected Stargardt disease (STGD) patients indicated that female sex could be a disease modifier [20]. *ABCA4* defects are clearly the most common IRD-causing mutations, with a predicted prevalence of 1.4 million globally [21]. In the context of RP, studies conducted in mouse disease models detected a more rapid loss of electroretinogram (ERG) responses in female rd10 mice [22] and accelerated retinal degeneration and vision loss in females of the P23H rhodopsin-mutant mouse strain [23,24]. Moreover, reserpine treatment promoted photoreceptor survival specifically in female P23H rats [25]. In addition, some aspects of the stronger retinal degeneration phenotype in females are also evident in mouse models of amyotrophic lateral sclerosis [26] and neuronal ceroid lipofuscinosis [27]. Collectively, the complex topic of sex differences in retinal degeneration requires further investigation with the best available disease models.

RP is in a privileged position from the translational perspective because many mutations cause very similar diseases across mammalian species, and many genetic defects underpinning the disease models occur spontaneously [28]. The translational potential of rd10 and P23H mouse models carrying mutations in the phosphodiesterase 6β (*Pde6b)* and rhodopsin (*Rho)* genes, respectively [29–31], is deemed to be particularly promising, leading to their high utilization in this research field. Here, we have carefully assessed the disease progression in both sexes of rd10 and P23H mice and observed that the rate of disease progression was accelerated in females *versus* males after the onset of puberty. We determined that the exacerbated disease in adult P23H females co-occurred with increased retinal lipid peroxidation, whereas inflammatory and total protein oxidation status was similar between sexes.

## Results

### Female retinitis pigmentosa mice undergo a faster retinal degeneration than males after puberty

To corroborate findings reported in the literature [22–24] and to provide independent assessment of the sex-related difference in rd10 and P23H RP mice, we assessed retinal function and morphology in several cohorts of mice by undertaking ERG recordings and optical coherence tomography (OCT) imaging longitudinally (Fig. 1, Supplemental Fig. S1-S5). First, rd10 mice housed in dim lighting were tested at postnatal ages P28, P42, and P56 (Supplemental Fig. S2). At P28, when postnatal retinal development is complete [32] but the eye is still growing slowly [33], female rd10 mice displayed stronger photopic ERG b-wave amplitudes (Supplemental Fig. 2A; RM ANOVA main effect of sex p = 0.0239, interaction p = 0.0008) and a thicker outer nuclear layer (ONL) (Supplemental Fig. 2B; main effect of sex p = 0.0436). At P42, when mice have reached puberty [34], the photopic ERG amplitude difference became reversed, such that it became stronger in males (Supplemental Fig. 2A; interaction p = 0.0018). At this point, no marked difference was evident in the analysis of the ONL thickness (Supplemental Fig. 2B). When male rd10 mice reached adulthood (P56), their ONL had become thicker (Supplemental Fig. 2B; main effect of sex p = 0.0067), indicative of a better photoreceptor survival, although there were no significant differences between the sexes in the amplitudes of the photopic ERG values.

**Figure 1.**
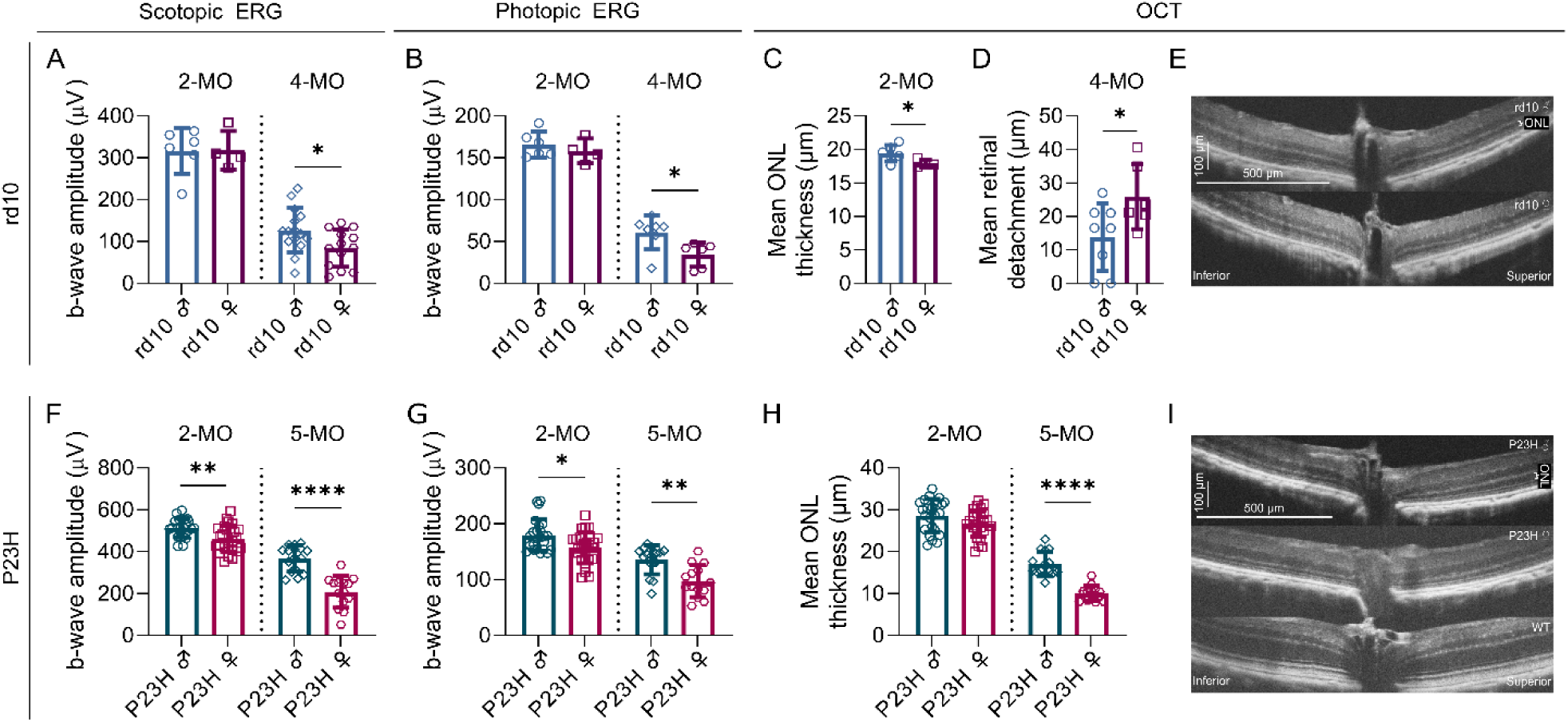
Functional and morphological sex difference in RP phenotype starts post-puberty and accelerates with aging in rd10 and P23H mice. Rd10 mice were constantly dark-reared whereas P23H mice were housed in standard vivarium conditions. A-B) B-wave amplitudes from scotopic (A; stimulus intensity: 30 cd • s/m2) and photopic (B; 50 cd • s/m2) electroretinogram (ERG) experiments in rd10 mice, mean ± SD. C) Mean outer nuclear layer (ONL) thickness from four retinal orientations (superior, inferior, nasal, temporal) in 2-month-old rd10 mice as measured with optical coherence tomography (OCT), mean ± SD. D) Mean retinal detachment (µm) from vertical retina in 4-month-old rd10 mice, mean ± SD. ONL thickness was not quantifiable from OCT images at this stage in rd10 mice. E) Example OCT images from 2-month-old male and female rd10 mice from the vertical retina. F-G) B-wave amplitudes from scotopic (F; stimulus intensity: 30 cd • s/m2) and photopic (G; 50 cd • s/m2) ERG experiments in P23H mice, mean ± SD. H) Mean ONL thickness from four retinal orientations as measured with OCT in 2- and 5-month-old P23H mice. I) Example OCT images from the vertical retina from 5-month-old male and female P23H, and healthy wild-type (WT) mice. The asterisks indicate significant Welch’s t-test (A, C-D, G, I) or Mann-Whitney test results (B, H), respectively: * = p < 0.05, ** = p < 0.01, **** = p < 0.0001. This figure presents a summary of phenotyping data. Comprehensive analyses are shown in Supplemental Fig. S2-S5.

Phenotypic variability in rd10 mice is a relatively well-known issue that is at least partially influenced by differing light exposure levels. These RP mice are particularly sensitive to light, even at vivarium conditions [35]. Additionally, the *Pde6b* mutation leads to unstable rather than fully absent enzyme activity causing variation in PDE6B levels and disease progression [36]. Indeed, our experiments with dim-light reared rd10 mice also led to high between-subject variability, regardless of sex. These mice were housed in a dimmed Scantainer® (Scanbur, < 1 lux), but still in a vivarium with a regular 12-hour light/dark cycle. To address the potential issue of varying environmental light exposure during housekeeping activities, we housed several litters of rd10 mice in complete darkness, where daily husbandry tasks were performed with only a very dim red light (> 600 nm wavelength). Two-month-old (∼P60) dark-reared rd10 mice did not show any sex-related differences in scotopic or photopic ERG amplitudes (Fig. 1A-B, Supplemental Fig. 3A, 3C). However, there was a marked sex difference in mean ONL thickness (Fig. 1C, 1E; Welch’s t-test p = 0.0321) as the rd10 female retina, especially at the superior retina (Supplemental Fig. 3E; main effect of sex p = 0.0072, interaction p = 0.0033), was thinner. When the disease had advanced to a more severe level in the 4-month-old (∼P120) dark-reared rd10 mice, the females had markedly lower scotopic (Fig. 1A; Welch’s t-test p = 0.0266; Supplemental Fig. 3A-B; main effect of sex p = 0.0008, interaction p = 0.0091) and photopic (Fig. 1B; Mann-Whitney test p = 0.0140; Supplemental Fig. 3C-D; main effect of sex p = 0.0160, interaction p < 0.0001) ERG b-wave amplitudes. By this age, the ONL was so thin that it was no longer possible to measure its thickness using OCT images. However, qualitatively, the retinal anatomy of the male mice appeared to be better preserved (Supplemental Fig. S4), and further, the females showed signs of a more severe retinal detachment (Fig. 1D; Welch’s t-test p = 0.0440; Supplemental Fig. S4; main effect of sex p = 0.0167).

Next, we examined P23H mice with a heterozygous mutation in the *Rho* gene. At 1 month of age, no sex differences were observed in either the functional (Supplemental Fig. 5A, 5C) or morphological (Supplemental Fig. 5E) RP phenotypes. At the age of 2 months (∼P60), P23H female mice showed lower scotopic (Fig. 1F; Welch’s t-test p = 0.0018; Supplemental Fig. 5A; main effect of sex p = 0.0005, interaction of sex and stimulus level p < 0.0001) and photopic ERG responses (Fig. 1G; Mann-Whitney test p = 0.0225; Supplemental Fig. 5C; main effect of sex p = 0.0325, interaction p = 0.0093) than their male littermates. However, at this time-point, there were no marked differences in ONL thickness (Fig. 1H, Supplemental Fig. 5E). The sex difference progressed between 2 and 5 months of age, such that the 5-month-old females exhibited markedly lower amplitudes in both scotopic (Fig. 1F; Welch’s t-test p < 0.0001; Supplemental Fig. 5A-B; main effect of sex p < 0.0001, interaction p < 0.0001) and photopic ERG recordings (Fig. 1G; Mann-Whitney test p = 0.0011; Supplemental Fig. 5C-D; main effect of sex p = 0.0008, interaction p < 0.0001). Furthermore, a marked structural difference was observed, as evidenced by the thinner ONL layer in female mice (Fig. 1H-I; Welch’s t-test p < 0.0001; Supplemental Fig. 5E-F; main effect of sex p < 0.0001, interaction p < 0.0001). As a control experiment, we also investigated 2- and 5-month-old C57BL/6J WT mice but did not detect any sex differences in either ERG responses or ONL thickness (Supplemental Fig. S6).

### Lipid peroxidation is specifically increased in adult female retinitis pigmentosa mice

The inflammatory response and oxidative stress are known to be closely linked with RP progression [37,38]. We hypothesized that either one or both factors could mechanistically be associated with the sex-related differences in the mouse RP phenotype. Initially, we tested the inflammatory status by performing immunoblotting of glial fibrillary acidic protein (GFAP) with 2- and 5-month-old WT and P23H retina samples. In the healthy state, GFAP is expressed in retinal Müller glial cells and astrocytes, but its levels dramatically increase if there is the presence of inflammation [37]. GFAP expression was markedly higher in both 2- and 5-month-old P23H retinas (Fig. 2A-B; 2-MO main ANOVA effect p = 0.0281, WT/P23H male comparison p = 0.0178, and WT/P23H female comparisons p = 0.0470; 5-MO main ANOVA effect p < 0.0001, WT/P23H male comparison p < 0.0001, and WT/P23H female comparisons p = 0.0003; Supplemental Fig. S7), but there were no sex-related differences at either of the time points. Next, we performed real-time quantitative polymerase chain reaction (qPCR) experiments and determined the expression of six typical inflammation marker genes: vimentin (*Vim*), integrin subunit alpha M (*Itgam*), interleukin 6 (*Il6)*, tumor necrosis factor alpha (*Tnfa*), interferon regulatory factor 8 (*Irf8*) and S100 calcium-binding protein B (*S100b*) with primer specificity being tested with electrophoresis after amplification (Supplemental Fig. S7). There was a clear effect of disease on the expression of *Vim* (Fig. 2C; main ANOVA effect p < 0.0001, WT/P23H male and WT/P23H female comparisons p < 0.0001) and *Itgam* (Fig. 2D; main ANOVA effect p < 0.0001, WT/P23H male and WT/P23H female comparisons p < 0.0001) in both 2- and 5-month-old P23H retinas. However, no sex differences were detected in any of the targets (Fig. 2C-H). Overall, it seems unlikely that differences in the inflammatory status could explain the sex difference in the P23H mouse RP phenotype.

**Figure 2.**
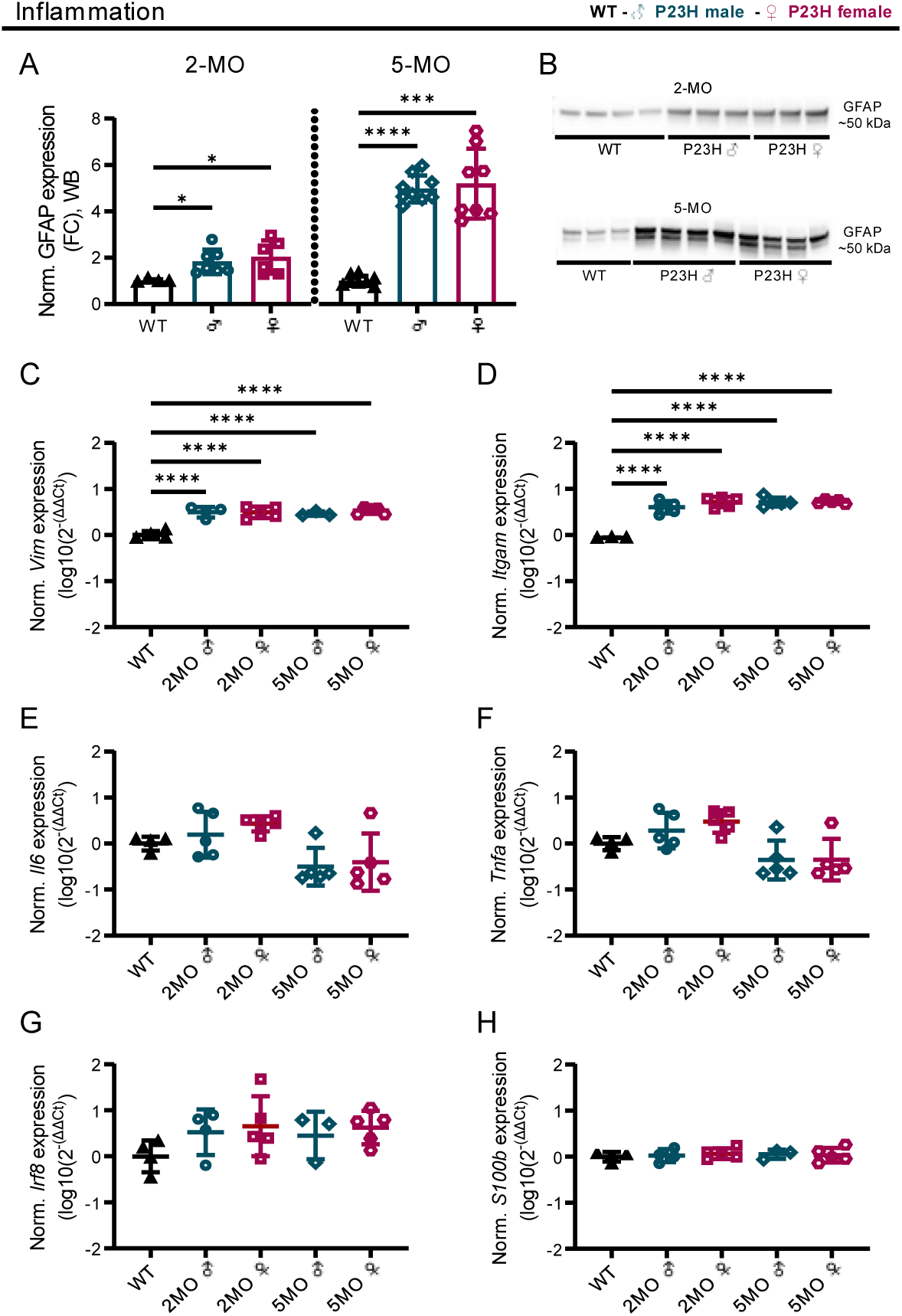
Inflammatory markers are increased by the same degree in male and female P23H mouse retinas. A) Normalized expression of glial fibrillary acidic protein (GFAP) in the immunoblotting/western blotting (WB) assay. Fold-change (FC) compared to WT, mean ± SD. B) Representative immunoblots as obtained from 2- and 5-month-old WT and P23H retina extracts. C-H) Expression of inflammatory marker genes C) vimentin (*Vim*); D) integrin subunit alpha M (*Itgam*); E) interleukin 6 (*Il6*); F) tumor necrosis factor alpha (*Tnfa*); G) interferon regulatory factor 8 (*Irf8*); and H) S100 calcium-binding protein B (*S100b*) as measured with real-time quantitative PCR (qPCR), mean ± SD. The asterisks indicate significant one-way ANOVA posthoc (Dunnett’s T3 used in panel A and Tukey’s test used in panels C-H) test results: * < 0.05, *** = p < 0.001 and **** = p < 0.0001.

To test the presence of oxidative stress, we performed quantitative measurements of standard markers of lipid peroxidation (4-hydroxynonenal, 4-HNE) and protein oxidation (protein carbonylation). Interestingly, the 4-HNE enzyme-linked immunosorbent assay (ELISA) detected an increased 4-HNE content in 1-month-old P23H male samples as compared to WT, whereas this was not the case in females (Fig. 3A; 1-MO main ANOVA effect p = 0.0157, WT/P23H male comparison p = 0.0140). However, after puberty, 4-HNE levels were elevated only in female P23H mice in comparison to WT mice, and furthermore, when the comparison was conducted between P23H female and male mice, this revealed a marked sex difference unfavorably for the females at both 2 and 5 months of age (Fig. 3A; 2-MO main ANOVA effect p = 0.0043, WT/P23H female comparison p = 0.0082, P23H male/P23H female comparison p = 0.0125; 5-MO main ANOVA effect p < 0.0001, WT/P23H female comparison p = 0.0001, P23H male/P23H female comparison p = 0.0009). The elevated 4-HNE expression was confirmed by immunohistochemistry (IHC) data (Fig. 3B-C; 2-MO main ANOVA effect p = 0.0315, WT/P23H female comparison p = 0.0255; 5-MO main ANOVA effect p = 0.0102, WT/P23H female comparison p = 0.0288, P23H male/P23H female comparison p = 0.0129; Supplemental Fig. S8). Next, we assessed the level of protein oxidation by measuring the total retinal protein carbonyl content with ELISA (Fig. 3D) and immunoblotting (Fig. 3E-F, Supplemental Fig. S9). Neither method revealed any significant evidence of sex- or disease-dependent changes in protein carbonyl levels. Finally, we assessed the retinal antioxidative system by first measuring the expression of key enzyme genes involved in clearing ROS and combating oxidative stress: superoxide dismutase 1 (*Sod1)* and 2 (*Sod2*), which are required to convert superoxide radicals into hydrogen peroxide, and catalase (*Cat*) and glutathione peroxidase 4 (*Gpx4*), which degrade hydrogen peroxide into water and oxygen [38]. Importantly, GPX4 is also required to combat the lipid peroxidation of PUFAs. As before, primer specificity was tested with electrophoresis after amplification (Supplemental Fig. S10). We did not find any significant differences in the expression levels of either *Sod1* (Fig. 3G) or *Sod2* (Fig. 3H) between WT and P23H retinas, regardless of sex. *Cat* was significantly overexpressed in P23H retinas compared to WT retinas, but there were no differences between P23H males *versus* P23H females (Fig. 3I; main ANOVA effect p = 0.0051, WT/2MO P23H male comparison p = 0.0068, WT/2MO P23H female comparison p = 0.0347, WT/5MO P23H male comparison p = 0.0268, WT/5MO P23H female comparison p = 0.0033). The expression of *Gpx4* was generally lower in P23H mouse retinas as compared to WT retinas, but this was statistically significant only between WT and 5-month-old P23H male mice (Fig. 3J; main ANOVA effect p = 0.0475, WT/5MO P23H male comparison p = 0.0346). In addition, we tested SOD2 and CAT protein expression with immunoblotting but found no sex differences (Fig. 3K-F, Supplemental Fig. S10). Lastly, the antioxidative function of the tissue was tested with a 2,2-diphenyl-1-picrylhydrazyl (DPPH) scavenging assay in retinas collected at 1 and 2 months of age, but no significant disease- or sex-dependent effects were observed in P23H mouse retinas (Fig. 3K).

**Figure 3.**
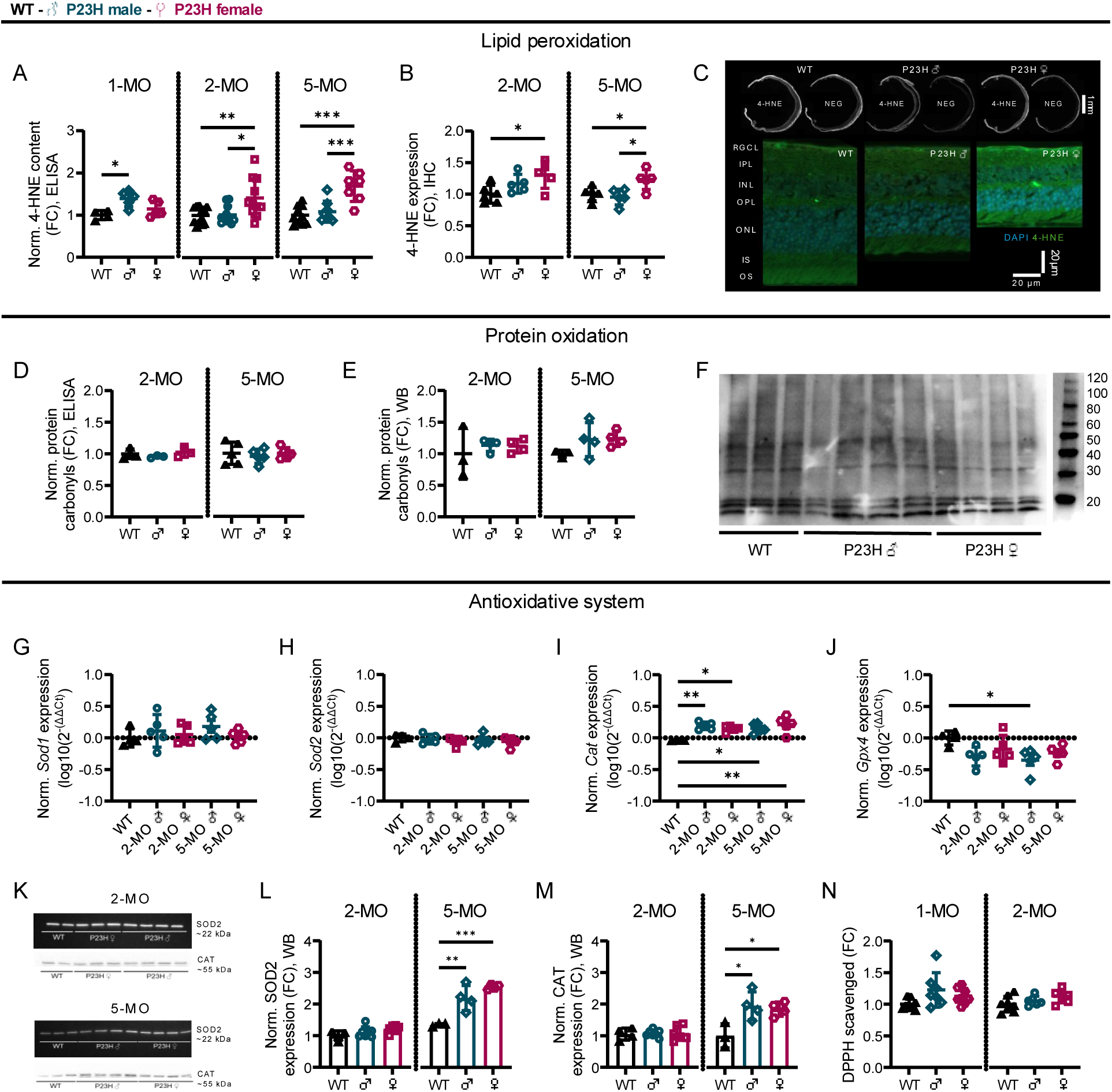
Significantly elevated retinal lipid peroxidation is evident in adult P23H female mice but not in P23H male mice. The figure presents normalized expression of A-F) oxidative stress markers, G-J) genes encoding reactive oxygen species (ROS) clearing enzymes, K-M) superoxide dismutase 2 (SOD2) and catalase (CAT) protein expression, and N) 2,2-diphenyl-1-picrylhydrazyl (DPPH) scavenging activity (a measure of antioxidant efficiency) across age and sex in P23H and WT mouse whole retinal extracts, mean ± SD. A) 4-hydroxynonenal (4-HNE) content as measured with an enzyme-linked immunosorbent assay (ELISA), fold-change (FC) to WT. B) 4-HNE expression as measured from antibody staining intensity in the immunohistochemistry (IHC) assay, FC compared to WT. C) Upper section: Representative raw images of 4-HNE signal used in the analysis with the corresponding negative control image from 5-month-old retinal cryosections – Lower section: Corresponding magnified IHC images as obtained from 5-month-old retinal cryosections stained with a 4-HNE antibody (green) and counterstained with the nuclei marker 4′,6-diamidino-2-phenylindole (DAPI; blue). D-E) Protein carbonyl content as measured with D) ELISA, or E) immunoblotting/western blotting (WB), FC compared to WT. F) A representative carbonyl immunoblot image presenting data obtained from 5-month-old mouse retinas. G-J) Expression of genes encoding ROS clearing enzymes G) *Sod1*; H) *Sod2*; I) *Cat*; and J) glutathione peroxidase 4 (*Gpx4*), as measured with quantitative polymerase chain reaction (qPCR), log10(2^-(ΔΔCt)^). K) Example images from SOD2 and CAT immunoblotting. L) SOD2 expression as measured with WB, FC to WT. M) CAT expression as measured with immunoblotting, FC to WT. N) DPPH scavenging activity, FC to WT. The asterisks indicate significant one-way ANOVA posthoc (Tukey’s) test results, * = p < 0.05, ** = p < 0.01 and *** = p < 0.001.

To increase the generalizability of the findings above, we analyzed the main parameters of inflammation and oxidative stress also in rd10 mouse retinas. We found significant elevation of 4-HNE content in 2-month-old rd10 female mice in comparison to WT and rd10 males (Fig. 4C; main ANOVA effect p = 0.0010, WT/rd10 female comparison p = 0.0007, rd10 male/rd10 female comparison p = 0.0467). No significant sex-dependent effects were observed in animals of 1 or 4 months of age. It is conceivable that by 4 months of age, retinal degeneration in rd10 mice is so advanced that a large proportion of 4-HNE–modified cells have already died off. Furthermore, there were no sex differences in GFAP expression (Fig. 4A-B, Supplemental Fig. S7), protein carbonylation (Fig. 4D), or DPPH scavenging activity (Fig. 4E), in accordance with the results from P23H mouse retinas. Curiously, DPPH scavenging activity was increased in both sexes of the rd10 mouse retinas in comparison to WT (Fig 4E; main ANOVA effect p = 0.0016, WT/1MO rd10 male comparison p = 0.0099, WT/1MO rd10 female comparison p = 0.0206, WT/2MO rd10 male comparison p = 0.0025, WT/2MO rd10 female comparison p = 0.0026). Collectively, it is conceivable that exacerbated lipid peroxidation could explain the more rapid disease progression in female RP mice after puberty; however, this is unlikely to be due to sex differences in antioxidant defense capacity.

**Figure 4.**
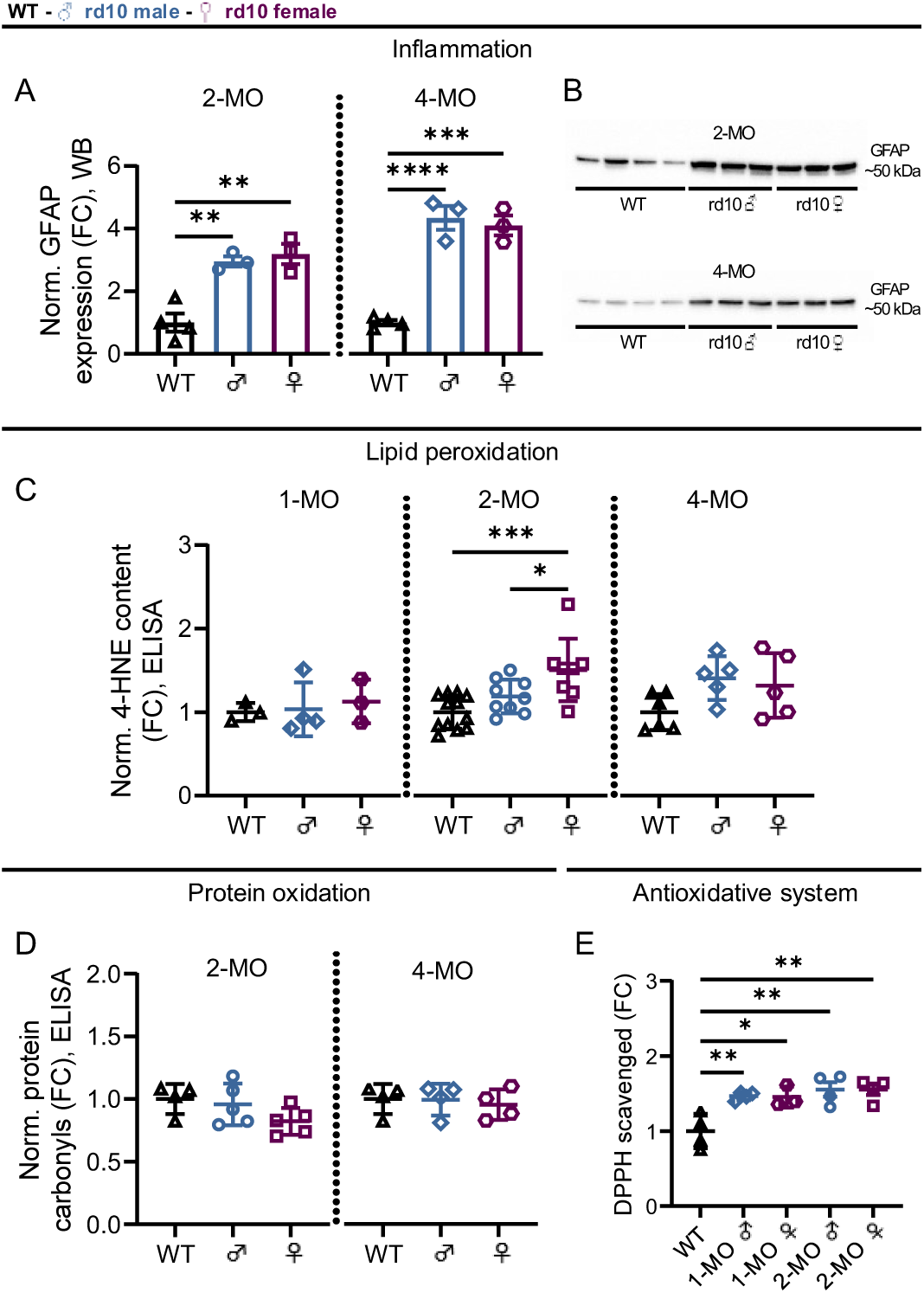
Investigation of inflammation and oxidative stress in rd10 retinas reveals higher lipid peroxidation in 2-month-old rd10 females, whereas inflammation and antioxidant capacity show only disease-dependent changes. A) Normalized expression of glial fibrillary acidic protein (GFAP) in the immunoblotting/western blotting (WB) assay, fold-change (FC) compared to WT. B) Representative immunoblots as obtained from 2- and 4 -month-old WT and rd10 retina extracts. C) 4-hydroxynonenal (4-HNE) content as measured with an enzyme-linked immunosorbent assay (ELISA), FC compared to WT. D) Protein carbonyl content measured with ELISA, FC compared to WT. E) Antioxidative capacity to scavenge 2,2-diphenyl-1-picrylhydrazyl (DPPH), FC to WT. Data is presented as mean ± SD, and the asterisks mark significant one-way ANOVA posthoc (Tukey’s) test results, * = p < 0.05, ** = p < 0.01, *** = p < 0.001 and **** = p < 0.0001.

### Global lipidomics demonstrate increased abundance of numerous lipid species in adult WT female retina relative to males

Given that lipid peroxidation was specifically increased in sexually mature RP female retinas, we next performed global lipidomics using liquid chromatography–mass spectrometry (LC-MS) in 1- and 2-month-old WT and P23H mouse retinas. After aligning the data, lipids with identification grade A or B (A: All class-specific ions and substituent-specific ions that specify the structure are assigned; B: Partial assignment with one class-specific ion and substituent-specific ion that specify the structure) were confidently assigned to a lipid class for further analysis. In 1-month-old WT male *versus* WT female and P23H male *versus* P23H female comparisons (Supplemental Fig. S11), lipid group classification A or B could be given to 30 and 47 differentially expressed lipids (DELs), respectively, and in both comparisons most of these lipids were triglycerides (TGs). When DELs belonging to the same lipid class were grouped together in the analysis (AUCs of lipids belonging to the same lipid class summed), retinas of 1-month-old WT females showed higher phosphatidylcholine (PC), TG, phosphatidylethanolamine (PE) and phosphatidylinositol (PI) content in contrast to WT male littermates. In the analysis of DELs between 1-month-old P23H males and females, females had higher abundance of lipids in PI class but lower PC, PE, phosphatidylglycerol (PG) and diglyceride (DG) lipid content. In post-pubertal retinas at 2 months of age, a total of 1024 lipids were detected (Fig. 5A), and 76 DELs could be confidently allocated to designated lipid classes (Fig. 5B). WT females showed higher abundance in all the lipid classes compared to WT males when DELs were grouped together (Fig. 5C). When all identified lipids (not just DELs) with lipid class assignment were included in the analysis, the difference remained significant with respect to PC and coenzyme Q (Co) lipid classes (Fig. 5D). Twenty-seven DELs in the 2-month-old P23H male and female comparison could be classified out of the 884 total compounds detected (Fig. 5E-F); sixteen of these were enriched in males, with most (9/16) representing PE lipids. The analysis of lipid groups based on DELs between P23H male and female groups detected a higher content of retinal PE, phosphatidylserine (PS), and PG in P23H males, whereas TGs were enriched in P23H females (Fig. 5G). However, no significant differences between male and female P23H retinas were present when the same analysis was performed with all classified lipids included (Fig. 5H).

**Figure 5.**
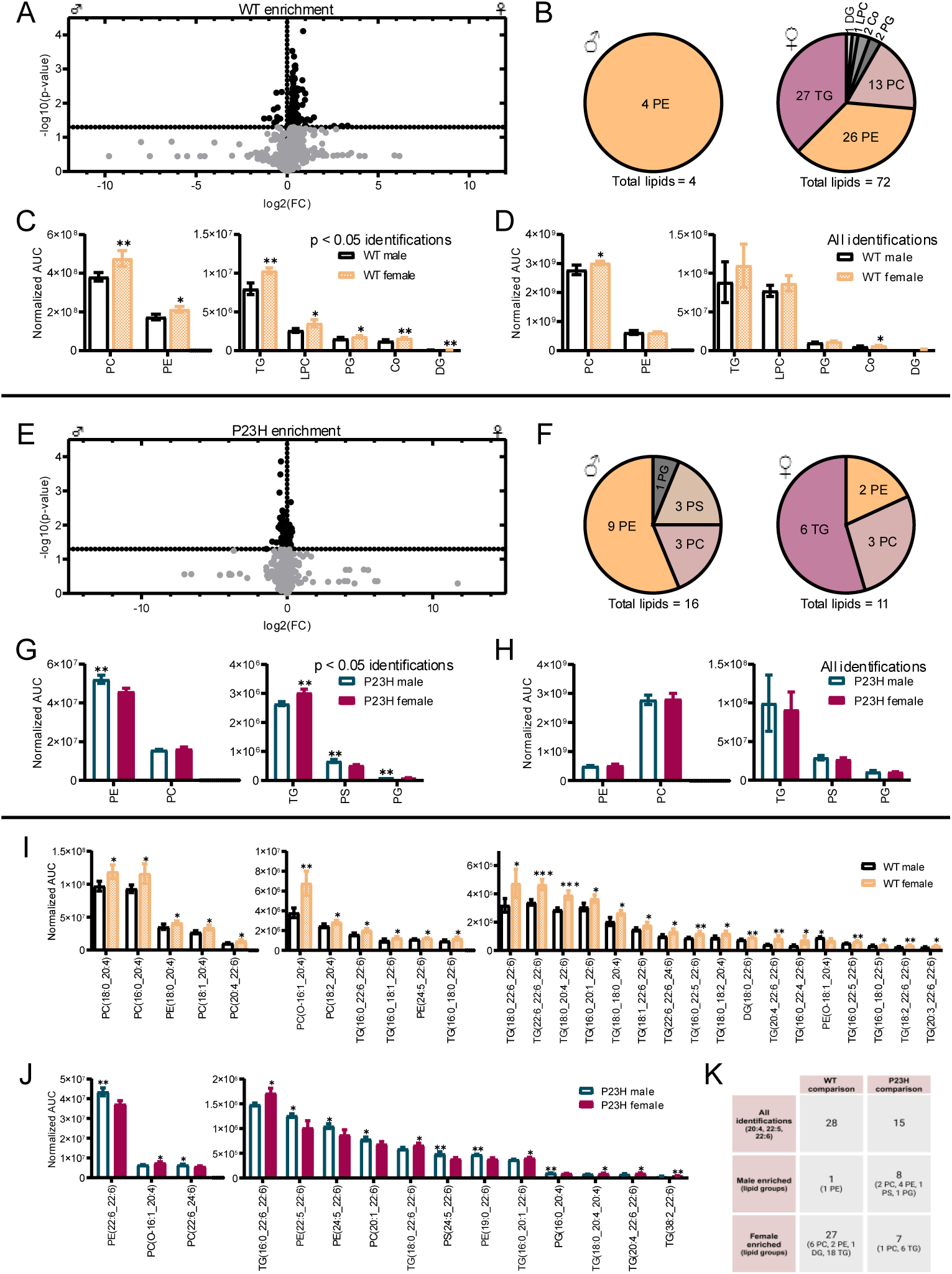
Global lipidomics demonstrate increased abundance of numerous lipid species in adult WT female retina relative to males. A) Volcano plot of all identified lipid species (regardless of identification confidence) in WT mice. Differentially expressed lipids (DELs, p < 0.05) between WT males (n = 4) and females (n = 4) are shown in black. B) Number of enriched grade A/B DELs per lipid class in WT females and males. C) Lipid group expression comparison between WT male and female mice based on grade A/B DELs only. D) Lipid group expression comparison between WT male and female mice based on all A/B lipid identifications. E) Volcano plot of all identified lipids in P23H mice (n = 3 m; n = 4 f). F) Number of enriched grade A/B DELs per lipid class in P23H females and males. G) Lipid group expression comparison between P23H male and female mice based on grade A/B DELs only. H) Lipid group expression comparison between P23H male and female mice based on all A/B lipid identifications. I) Grade A DELs with 20:4/22:5/22:6 carbon/double-bond sidechains (grade A) between WT male and female mice, and J) between P23H male and female mice. K) Summary of lipid group distributions based on DELs with 20:4/22:5/22:6 carbon/double-bond sidechains in WT and P23H comparisons. In all graphs, the asterisks indicate significant t-test results, * = p < 0.05, ** = p < 0.01 and *** = p < 0.001. The bar graph data is presented as mean ± SD. Co = coenzyme Q, DG = diglyceride, LPC = lysophosphatidylcholine, PC = phosphatidylcholine, PE = phosphatidylethanolamines, PG = phosphatidylglycerol, PS = phosphatidylserine, TG = triglyceride.

Long-chain PUFAs (LC-PUFAs) are particularly important for the normal structure and function of the retina [39]. However, these compounds are also susceptible to oxidative damage due to their chemical structure. In the comparison of DELs between 2-month-old WT female and male retinas, we identified 28 molecules (identification grade A) with 20:4, 22:5, or 22:6 carbon/double bond sidechains, which are markers for LC-PUFAs *e.g.*, arachidonic acid (AA), docosapentaenoic acid (DPA), and docosahexaenoic acid (DHA), respectively [39,40]. Apart from one LC-PUFA-containing lipid, the remaining (27/28) were significantly more abundant in the female retinas (Fig. 5I, 5K). When all grade A identifications with these sidechains (not just DELs) were considered, a higher prevalence of PC and TG lipids with LC-PUFA sidechains persisted in female WT retinas (Supplemental Fig. S12). In contrast, such female-specific enrichment of LC-PUFAs was absent in 1-month-old WT retinas: only seven DELs with 20:4, 22:5, or 22:6 sidechains were detected, and these were more evenly distributed between sexes (three enriched in males and four in females, Supplemental Fig. S11). In the comparison of P23H male *versus* P23H female retinas at 2 months, 15 DELs with 20:4, 22:5, or 22:6 carbon/double bond sidechains were detected, of which 8 were enriched in males and 7 in females (Fig. 5J). Interestingly, 18 out of the 21 DELs with LC-PUFA sidechains were elevated in 1-month-old P23H male retinas in contrast to the female tissue (Supplemental Fig. S11). In both WT-WT and P23H-P23H comparisons with adult retinas, TGs represented most of the female-enriched lipids with LC-PUFA sidechains (Fig. 5I-K). In addition, TGs appeared to be particularly altered due to the disease, as differentially expressed TGs with or without LC-PUFA sidechains were elevated in the P23H retinas regardless of sex or age (Supplemental Fig. S13-S14). Furthermore, the disease-effect on LC-PUFA containing TGs was significant in 2-month-old mice when all grade A lipids with sidechains were included in the analysis (Supplemental Fig. S12).

Collectively, these findings indicate that after puberty, WT female retinas accumulate significantly more LC-PUFA–containing lipids than males, potentially increasing lipid peroxidation risk.

### Open transcriptomic data suggests sex differences in energy metabolism in middle-aged human retinas

Human retinal lipidomic data to evaluate sex-related differences is not readily available since it is extremely challenging to obtain this data from middle-aged or younger subjects, particularly for performing well-powered differential expression analyses. As surrogate data, we downloaded and re-analyzed open retinal bulk RNA-seq data [41] obtained from ocularly healthy subjects between the ages 42 and 59 (mean SD of age of females 53.7 ± 4.4, n = 7, and males 51.1 ± 4.4, n = 10; p = 0.36), excluding samples where cause of death was marked as “cardiovascular disease”. We identified 32444 autosomal genes in our analysis (Fig. 6A), and 66 of them reached the statistically significant criterion (adjusted p-value/q < 0.1) of being differentially expressed genes (DEGs) between sexes. The retinal DEGs enriched in females included 22 genes, such as dipeptidyl peptidase 4 (*DDP4*), which is involved in the regulation of glucose metabolism [42], and acetyl-CoA acyltransferase 2 (*ACAA2*), which encodes for the enzyme involved in the final step of the β-oxidation of fatty acids [43]. DEGs enriched in male retinas included the gluconeogenesis regulator phosphoenolpyruvate carboxykinase 1 (*PCK1*) [44], transcription regulators FosB proto-oncogene (*FOSB*) [45] and early growth response proteins (*EGR*) 1, 2 and 3 [46], and *PDE10A*, which eliminates cyclic nucleotides [47]. Heatmaps of all genes (no statistical criteria applied) involved in the KEGG pathways “glycolysis/gluconeogenesis” (Fig. 6B), “citric acid cycle” (Fig. 6C) and “fatty acid metabolism” (Fig. 6D) provide further insights into the possible sex differences in retinal metabolism. Taken together, although the sample size here is small and definitive roles of physiological functions cannot be allocated solely from gene expression data, this analysis does provide an initial indication that human female and male retinas may differ in terms of their energy metabolism pathways.

**Figure 6.**
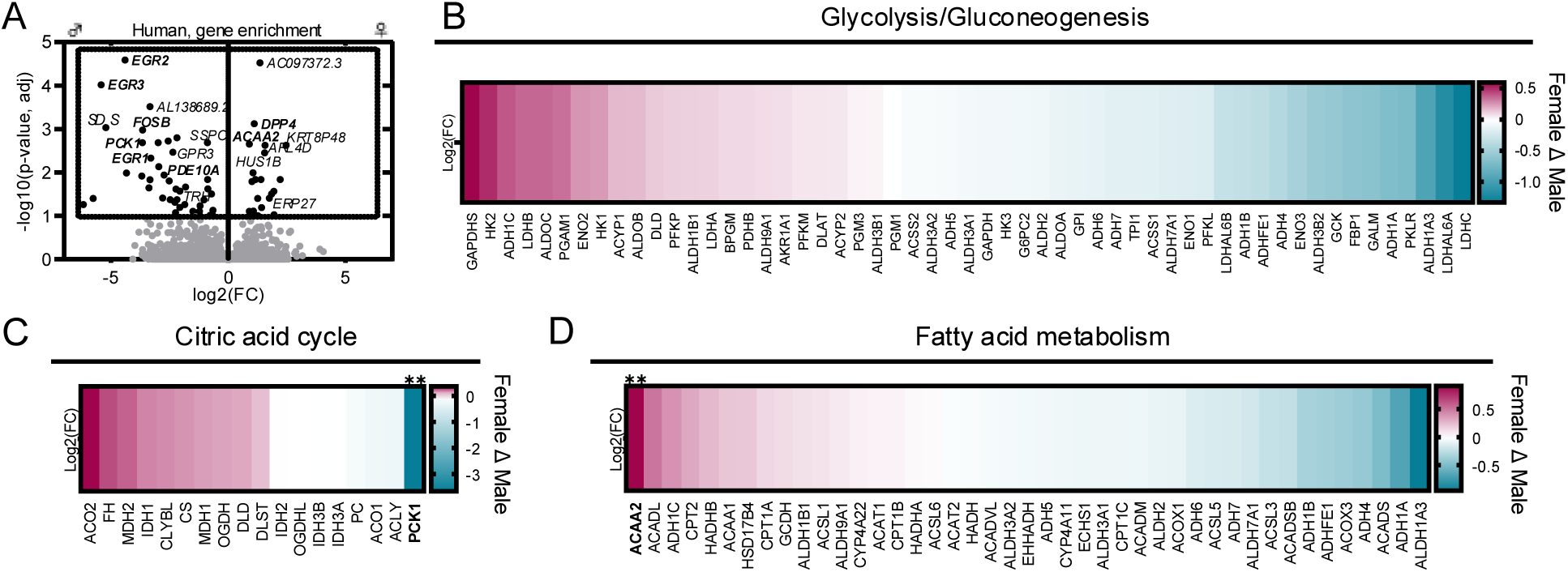
Analysis of open retinal transcriptomics data suggests the presence of sex differences in mechanisms of energy production in middle-aged humans. A) Volcano plot of all identified autosomal genes. Differentially expressed autosomal genes (DEGs, adjusted p-value/q < 0.1) between males (n = 10) and females (n = 7) are shown in black. Heatmaps of all genes (no statistical criteria applied) that belong to the KEGG pathways for B) glycolysis/gluconeogenesis, C) citric acid cycle, and D) fatty acid metabolism. DEGs between sexes in heatmaps are bolded and \marked with asterisks, ** = q < 0.01.

## Discussion

RP is the most prevalent form of the IRDs, affecting approximately 1/4000 individuals globally [48], and sadly, no broadly effective treatment is yet available. RP is characterized by rod degeneration with a periphery-to-central field progression gradient, leading to the loss of night vision and visual fields as the first clinical manifestations of the disease. Collateral cone degeneration eventually occurs impacting on color and high-acuity vision [1,48]. Sex differences in predisposition, age of onset, or progression, have not been reported in autosomal recessive, autosomal dominant or rare (mitochondrial, digenic, sporadic) disease types. The X-linked RP, caused by mutations in the X chromosome, affects males more commonly whereas females are carriers and suffer only mild or no symptoms. However, the high genetic heterogeneity and rarity of individual RP subtypes have significantly hindered the identification of disease modifiers in this disease group.

In accordance with previous results [23,24], our follow-up of the phenotype in RP mice indicates that the disease accelerates in females after the animals have reached puberty (Fig. 1, Supplemental Fig. S2-S5). In adult mice (P>60) the phenotype was consistently aggravated in females of both rd10 and P23H RP mouse models, and in P23H mice at 5 months of age, the sex difference in retinal function could be as high as 50 % (Supplemental Fig. S5A).

We investigated retinal inflammation and oxidative stress as potential contributors to the observed functional and morphological sex differences in RP mice. The main focus of our investigations was on P23H mice, owing to the lower inter-subject variability in this model and the higher prevalence of *RHO* mutations compared to *PDE6B* mutations in humans [48]. Nevertheless, results from complimentary experiments with rd10 retinas were in accordance with the P23H experiments (Fig. 4). Our immunoblotting and qPCR analyses with WT and P23H retinas (Fig. 2) revealed that GFAP protein and *Vim* gene expressions were equally increased in female and male P23H mouse retinas, confirming findings that the level of Müller cell gliosis is similar between the sexes [49]. Retinal *Itgam* expression was elevated in P23H mice, indicating microglial activation, but this effect was not sex-dependent. Together with the absence of significant differences in *Il6*, *Tnfa*, *Irf8*, and *S100b* expression between WT and P23H retinas, these results suggest that retinal inflammation in P23H mice is characterized primarily by Müller cell gliosis and resident microglial activation, whereas astrocytosis or macrophage infiltration appears less pronounced or absent [49]. In contrast to our current results obtained from relatively young mice, sex differences in inflammatory gene expression have been observed in aged mouse retinal pigment epithelium (RPE) [50]. Moreover, eye transcriptomics in aging humans has revealed a sex-specific enrichment of inflammatory response genes in female retinal glial cells [51]. The levels of several inflammation markers, including the expression of *Il6* and *Tfna*, have also been shown to be elevated in female rd10 mouse and RCS rat retinas in contrast to males [51]. However, differently from the settings we used here, the animals examined in the study of Liu et al. (2025) [51] had been light-reared, meaning that the disease progression in the rd10 mice would have been significantly more aggressive [35]. The retinas of RCS rats were tested at P60, at which time they would have also been more severely degenerated than in our P23H mouse model at the age of 5 months [52]. Thus, while the signs of inflammation have displayed female-specific increases in aging or later stage RP retinas, this seems to be more likely a consequence of the disease rather than a driver.

Instead of inflammation, we argue that lipid peroxidation can provide a better explanation for the sex differences evident in the RP mice. It is known that excessive lipid peroxidation by ROS generates a detrimental chain reaction which disrupts lipid membrane integrity and induces cell death [38]. We detected increased levels of lipid peroxidation in 2-month-old female P23H mouse retinas in comparison to WT retinas as measured by ELISA and IHC (Fig. 3A-C). The same phenomenon was not observed in P23H males. This gap between the sexes became even more exacerbated by 5 months of age. In contrast, we did not find any significant elevation in retinal total protein oxidation levels in either female or male P23H mice (Fig. 3D-F). Overall, the literature on lipid peroxidation in RP is rather comprehensive [53–55], while protein oxidation is far less reported.

Next, we examined the retinal antioxidative system to test if it could contribute to the observed sex difference in lipid peroxidation. It has been reported that the expression levels of *Sod1* and *Sod2* are elevated in 8-month-old male mice with experimental AMD, a model based on a deficiency of peroxisome proliferator-activated receptor gamma coactivator 1-alpha (PGC-1α) [56]. Therefore, we tested the gene expression of these crucial antioxidative enzymes, as well as *Cat* and *Gxp4,* but did not detect any significant differences between P23H male and female retinas (Fig. 3G-J). In addition, we assessed the expression of SOD2 and CAT proteins with immunoblotting and tested antioxidative capacity using the DPPH scavenging assay, and again, found no differences between P23H male and P23H female retinas (Fig. 3K-N). Together, these results indicate that sex-dependent differences in retinal antioxidant defenses are unlikely to account for the increased lipid peroxidation observed in female RP mice.

Because the susceptibility of lipids to oxidative damage is largely determined by their composition, we next performed global lipidomics to assess whether sex-related differences in the retinal lipid pool might contribute to this phenomenon. Our analysis from 2-month-old sexually mature mice revealed that healthy WT female retinas were enriched in a broader range of lipids than male retinas, with LC-PUFA-containing species especially prominent (Fig. 5A-D, 5I, 5K). Instead, there was a greater disease-related lipid downregulation in P23H females than in males (Fig. 5E-G, Supplemental Fig. 14). The retina is particularly susceptible to oxidative damage by lipid peroxidation, as the tissue is highly enriched in various lipids, particularly LC-PUFAs, which are typically found within TGs or phospholipid structures such as PCs [39]. While also crucial for healthy retina function, excessive amount of LC-PUFA DHA (22:6) is known to exacerbate lipid peroxidation in RPE cells [57]. Our lipidomics analysis of prepubertal (1-month-old) WT retinas did not reveal significant sex-dependent enrichment of LC-PUFA-containing lipids (Supplemental Fig. S11).

One possible explanation for the female-specific increase in lipid peroxidation we observed is the effect of sex hormones on retinal lipid composition. LC-PUFAs, particularly DHA, are generally more abundant in female tissues than in males [58–62], a phenomenon attributed to estrogen-driven synthesis, while testosterone treatment decreases LC-PUFA content [58]. In line with this, Rowe *et al.* [24] recently demonstrated reported that the sex difference in the P23H mouse phenotype can be counteracted by depleting sex hormones through ovariectomy. They further showed that estradiol plays a particularly important role, as its exogenous supplementation aggravated the retinal degeneration phenotype in ovariectomized female P23H mice. In contrast to results from adult RP mouse retinas, photopic ERG responses were *lower* in juvenile male rd10 mice (∼P28) reared in dim-lighting conditions (Supplemental Fig. S2). Second, we detected a modest *increase* in 4-HNE content in 1-month-old P23H male retinas compared to WT, which was not yet evident in females at this age (Fig. 3A). An important consideration that could explain the bidirectionality of disease parameters in young *versus* adult mice is that the earliest experiments at P28-P30 were performed at a stage when mice, including the size of their eyes, are still growing [63]. We speculate that differences in the speed of postnatal development or metabolism between the sexes could play a role in the sex-related findings that are present before puberty, when major differences in sex hormones males and females have not yet emerged [24,64].

It is well established that male and female tissues, including the retina [65], differ in terms of metabolic pathways due to the actions of the sex hormones [66,67]. Our results show markedly higher expressions of coenzyme Q9 and Q10 in 2-month-old WT female retinas in contrast to healthy males (Fig. 5C-D, Supplemental Data S02) which could reflect differences in oxidative phosphorylation capabilities [68]. While glucose is the main energy source of the retinal cells, photoreceptors can additionally utilize fatty acids for energy production via β-oxidation [69,70]. Importantly, our mouse retina data demonstrated sex- and disease-dependent upregulation particularly in TGs with LC-PUFA sidechains (Fig. 5I-K, Supplemental Fig. S12-S14). β-oxidation of fatty acids, including LC-PUFAs which are typically released specifically from TGs, occurs in peroxisomes and mitochondria after conjugation with coenzyme A (CoA) or carnitine, respectively. It has been reported that female WT mouse retina, brain, and plasma are enriched in the CoA precursor, pantothenic acid, and in the fasted state, the female retina has a higher long-chain acylcarnitine content [65], which may be evidence of a higher fatty acid oxidation rate in females. In line with this proposal, our analysis of open bulk retina RNA-seq data from middle-aged humans revealed elevated expression of *ACAA2* in females and suggests sex-related differences in the energy metabolism pathways in general (Fig. 6). Although the literature on neural and retinal lipid metabolism is limited and requires more attention, it has been reported that the metabolism of fatty acids is greater in female skeletal muscle and cardiac tissue, which has been speculated to be linked to their estrogen-induced elevated fatty acid content and higher level of β-oxidation due to higher enzyme availability [71–75]. If the same holds true for the retina, higher lipid turnover via β-oxidation could pose an additional burden on the female retina via lipid-related oxidative stress, especially in the presence of a degenerative disease where energy production mechanisms are dysregulated [69,70]. Given that female sex hormones additionally influence the progression of retinal degeneration [24], further investigations on the possible connection between retinal metabolism and sex hormones in RP is warranted. One interesting pathway worth consideration is that involving mitochondrial PGC-1α, as its inhibition has been reported to evoke RPE lipid droplet accumulation and lipid peroxidation *in vitro* [76] and it was claimed to account for the sex differences in retinal function in an *in vivo* model of AMD [56]. Furthermore, supporting proteomic data about sex differences in the retina could guide future investigations about the topic [77].

In conclusion, our findings show that post-pubertal female RP mice exhibit accelerated retinal degeneration that coincides with increased lipid peroxidation. This sex difference was not explained by inflammation or antioxidant capacity. These results underscore the importance of considering sex as a biological variable in retinal research.

## Materials and methods

### Animals

Heterozygote P23H (Rho^P23H/WT^, RRID: IMSR_JAX:017628), homozygote rd10 (Pde6β^R560C/R560C^, RRID: IMSR_JAX:004297), and WT C57B6/6L (RRID: IMSR_JAX:000664) mice were used in this study. P23H and WT mice were bred and housed in the Laboratory Animal Center, Kuopio, Finland, in a controlled environment (temperature 21 ± 1 °C, humidity 50–60 %, light period 07:00-19:00). The first cohorts of rd10 mice (P28, P42, P56) were housed in a fully dark ventilated cabinet in the same regular animal housing room as P23H and WT mice (1-, 2- and 5-month-old) and handled under dim red light and only exposed to dim background light during husbandry duties. Later rd10 cohorts (2- and 4-month-old) were fully dark-reared and handled under dim red light. Food and water were available *ad libitum*. The experiments were conducted according to the Council of Europe (Directive 86/609) and Finnish guidelines and approved by the Animal Experiment Board of Finland.

### Electroretinography

Retinal function was assessed with scotopic and photopic ERG. In the scotopic ERG experiments, rodents were dark-adapted overnight, and the pupils were dilated with a 1:5 mix of metaoxedrin and tropicamide (Oftan metaoksedrin 100 mg/ml, Oftan Tropicamid 5 mg/ml, Santen Oy) under dim red lighting. Mice were anesthetized with medetomidine (Domitor Vet 7.5 mg/ml, Orion Pharma) and ketamine Ketaminol Vet 0.1 mg/ml, MSD) solution in 0.9 % saline with a dose of 0.1 ml per 10 g of bodyweight.

A custom scotopic ERG protocol was utilized with the Espion E3 system and a Ganzfeld stimulator Colordome (V6.64.9, Diagnosys LLC), with the animal resting on a heating pad at 37.0 °C (Physiological-Biological Temperature Controller TMP-5b, Supertech). The reference and ground electrodes were placed under the skin of the snout, and the recording silver-plated electrodes were placed on the carbomer gel (Viscotears 2 mg/g, Novartis) coated corneas. The photopic ERG protocol was performed in light-adapted animals using an Espion Celeris rodent ERG device (D430, Diagnosys LLC). A rod-suppressing white background light at 20 cd • s/m^2^ was used. The full ERG protocols are listed in the Supplemental Tables S1 and S2. The ERG signal was acquired at 2 kHz and filtered with a low-frequency cutoff at 0.25 Hz and a high-frequency cutoff at 300 Hz. The Espion software automatically detected the ERG a-wave (first negative ERG component) and b-wave (first positive ERG component) amplitudes; the a-wave amplitude was measured from the signal baseline, whereas the b-wave amplitude was assessed as the difference between the negative trough (a-wave) and the highest positive peak. Only b-wave amplitudes were used in the statistical analysis.

### Optical coherence tomography

Retinal OCT b-scan images were captured with Phoenix Micron IV system (Phoenix-Micron) after ERG. Pupil dilation and anesthesia were performed as described above, and the corneas were lubricated with carbomer gel. Cross-sections of the retinal layers were rendered from vertical and horizontal orientations centered on the optic nerve. The thickness of the ONL was measured and averaged at 500 µm from the optic nerve head from all retinal quadrants (nasal, temporal, superior and inferior) by using an ImageJ ruler tool (Version 1.53k, National Institutes of Health).

### Quantitative real-time polymerase chain reaction

Mice were sacrificed under anesthesia with cervical dislocation, with retinas being immediately dissected from the eyecup and snap-frozen prior to RNA extraction. The samples were stored at −80 °C and soaked overnight in RNAlater-ICE solution (AM7030, Ambion) in −20 °C prior to processing. RNA extraction was conducted with an RNeasy Mini Kit (74104, Qiagen) according to the manufacturer’s manual with the addition of on-column DNase digestion with RNase-Free DNase Set (79254, Qiagen). RNA concentrations and quality were measured with Denovix DS-11, and RNA concentrations were equalized with water for molecular biology (BN-51100, BioNordika). The synthesis of cDNA was carried out with iScript cDNA Synthesis Kit (1708891, Bio-Rad) according to the manufacturer’s instructions, and the final product was diluted 5X with water. For qPCR, a reaction mixture of PowerUp SYBR Green Master Mix (A25741, Applied Biosystems, 5 µl per sample), forward and reverse primers (Supplemental Table S3, 0,5 µl per sample per primer) and water (3 µl per sample) was prepared and 9 µl of the final solution was added per well to a PCR 96 well-plate (732-2389, VWR). The samples were tested as three replicates with a sample volume of 1 µl per well. The well-plate was covered (89087-690, VWR) and centrifuged briefly to remove any air bubbles, and the PCR reaction was initiated and measured with QuantStudio 5 qPCR device (Applied Biosystems) in 3 stages (Stage 1: 2 min at 50 °C, 2 min at 95 °C; Stage 2: 15 s at 95 °C, 30 s at 60 °C, repeated 40 times; Stage 3: 15 s at 95 °C, continuous 1 s at 95 °C). The results were analyzed with Microsoft Excel and calculated with the ΔΔCt method by using succinate dehydrogenase complex flavoprotein subunit A (*Sdha*) and glyceraldehyde 3-phosphate dehydrogenase (*Gapdh*) expression for normalization and WT group as the control group.

### Immunohistochemistry

Mice were sacrificed and the eyes were enucleated. The corneas were cut, and the remaining eye was fixed in 4 % (V/V) paraformaldehyde (15710, Electron Microscopy Sciences) for 1 h, after which the iris and the lens were removed. Eye cups were dehydrated in sucrose solutions (S9378, Sigma-Aldrich, 10%/20%/30% (m/V) in 1XPBS), equilibrated overnight in the embedding media, embedded on the next day in 1:2 20%-sucrose/Frozen Section Media (3801480, Leica Biosystems), frozen on a liquid nitrogen-cooled metal block and stored at −80 °C. Cryosections were collected with Leica Cryostat CM3030 S to microscope slides (J1800AMNZ, Epredia) with a sample thickness of 10 µm.

Antibody staining was performed by first surrounding the sample with a hydrophobic barrier (H-4000, ImmEdge) and washing away the embedding media with 1XPBS for 10 min; this was followed with a 30 min incubation with a 10 % (V/V) normal donkey serum (NDS, Biowest) blocking solution, and with an overnight primary antibody in PBST (0.1 % (V/V) Tween 20) with 1 % NDS (anti-mouse 4-HNE 1:500, MAB3249-SP, R&D Systems) incubation at 4 °C. On the next day, the samples were incubated for 2 h with the secondary antibody (donkey anti-mouse Alexa Fluor 488 1:1000, AB150109, Abcam) at room temperature (RT) in the dark after washing with PBST for 5 min 3 times. After the incubation with the secondary antibody, the sections were again washed 3 times with PBST, twice with PBS and finally covered with a 4′,6-diamidino-2-phenylindole (DAPI) containing mounting medium (AB104139, Abcam). The samples were imaged with Leica Thunder Imager 3D Tissue Slide Scanner and the mean grey area of the whole retina was measured with ImageJ (Version 1.53k, National Institutes of Health). The results were normalized to the mean of WT group.

### Immunoblotting

Retina samples for immunoblotting were collected by dissection after sacrificing the mice with CO_2_. The samples were snap-frozen in liquid nitrogen and stored at −80 °C. Single retina samples were incubated for 10 min in 60 µl of ice-cold 1XRIPA (89900, Thermo Scientific) buffer with 1 % protease inhibitor (cOmplete™ Mini EDTA-free Protease Inhibitor Cocktail, 11836170001, Roche) and homogenized with a handheld device. After a further 15 min incubation on an ice bed, the samples were centrifuged at 4 °C for 20 min at 12 000 rpm, and sample and protein assay aliquots were taken. The sample volume was doubled with 2XLaemli buffer (1610737, Bio-Rad) with β-mercaptoethanol (β-ME, M3148, Sigma-Aldrich), and the samples were boiled at 95 °C for 5 min and cooled on ice for 5 min. Protein concentrations were equilibrated according to the Pierce BCA protein assay (23250, Thermo Scientific), which was conducted as per the manufacturer’s instructions. Gel electrophoresis was done with precast 4-20 % gels (4568095, Bio-Rad) and run at 200 V for 30 min. Gels were transferred to PVDF membranes (Trans-Blot Turbo Midi 0.2 µm PVDF Transfer Packs, 1704157, Bio-Rad) with Bio-Rad Trans-blot Turbo System and blocked in 5 % (m/V) fat-free milk (Skimmed milk powder, Valio) in TBST (with 0.1 % Tween 20, BP337-500, Fisher BioReagents) for 1 h at RT. After 3 washes with TBST, the membranes were incubated overnight in primary antibody (anti-mouse GFAP 1:2000 (3670, Cell Signaling Technology), anti-rabbit SOD2 1:5000 (ab13533, Abcam), anti-mouse CAT 1:500 (sc-271803, Santa Cruz Biotechnology)) in 5 % milk TBST. On the next day, the membranes were washed again with TBST and placed in the secondary antibody (anti-mouse HRP 1:5000 (A10668, Invitrogen), anti-rabbit IRDye 800CW 1:10000 (925-32211, Licor)) in 5 % milk TBST, and incubated for 2 h at RT. Imaging was done with Azure Biosystems 600 after washes with TBST and finally TBS. Chemiluminescence was induced with SuperSignal West Pico PLUS (34579, Thermo Scientific) according to the product’s protocol. Results were analyzed with ImageJ blot tool (Version 1.53k, National Institutes of Health) and normalized to the total protein signal assessed with Ponceau S (A40000278, Thermo Scientific) staining according to the manufacturer’s instructions and to the mean of WT group.

### 4-HNE ELISA

Retina samples were collected fresh after sacrificing the animal with CO_2_ and cervical dislocation, snap-frozen and stored at −80 °C. Samples were immersed in 110 µl of ice-cold 1XPBS (pH 7.4) with 1 % (V/V) protease inhibitor, homogenized with a plastic pestle, sonicated, and centrifuged for 10 min at 5000 G at 4 °C. 4-HNE ELISA (E-EL-0128, Elabscience) was conducted according to the manufacturer’s instructions.

### Protein carbonylation ELISA

Protein carbonylation was tested with a modified ELISA method [78]. Single retina samples were homogenized by sonication in ice-cold 1XPBS (pH 7.4) with 1 % protease inhibitor and 0.05 % (V/V) Triton-X (648466, Calbiochem) in a volume of 25X tissue weight (1 mg = 25 µl), and centrifuged for 10 min at 10000 G at 4 °C. The protein content of the supernatant was measured with the Pierce BCA protein assay (23250, Thermo Scientific), and nucleic acids were removed by incubating the supernatant in RT for 15 min with 1 % (V/V) streptomycin (85884, Sigma-Aldrich) and by collecting the supernatant after centrifuging for 5 min at 10000 G at 4 °C. Oxidized bovine serum albumin (BSA) stock (50 mg/ml) was produced by the Fenton reaction by incubating with BSA (A2153, Sigma-Aldrich), 1 mM FeCl_2_ (44939, Sigma-Aldrich), 1 mM EDTA (20301.290, VWR Chemicals) and 2 mM H_2_O_2_ (H1222, TCI) for 3 h at 37 °C and filtering the product with a Microcon-30kDa Centrifugal Filter Unit with an Ultracel-30 membrane (MRCF0R030, Millipore). The protein concentration of the 10X diluted solution was measured with the protein assay and serial dilutions (2, 1.5, 1,25, 1, 0.5, 0.1, 0.02 mg/ml) in water were made according to the value obtained. To generate a reduced BSA stock solution (10 mg/ml), BSA and sodium borohydride (16940-66-2, Merck) were incubated together for 30 min in 1XPBS at RT, neutralized to pH 7 and dialyzed overnight in 1XPBS. The protein concentration of the 10X diluted stock was assessed by measuring absorbance at 280 nm, and a 1 mg/ml solution was made based on the result by diluting with 1XPBS. In the assay, 15 µl of each standard, blank (lysis buffer), and the sample were incubated with 45 µl of 2,4-dinitrophenyl(hydrazine) (DNP, A5531, TCI) solution (10 mmol of DNP in 6 mol guanidine hydrochloride (G3272, Sigma-Aldrich) in 0.5 M potassium phosphate buffer, pH 2.5) for 45 min at RT. The solution was diluted by adding 10 µl to a tube with 2 ml of 1XPBS, mixed well and incubated overnight at 4 °C in Nunc Immuno Plate Maxisorp well plates (442404, Thermo Scientific) as three triplicates (50 µl/well). On the next day, the plate was washed 5 times with PBST (0.05 % Tween 20, 200 µl/well) and the wells were blocked with 200 µl of blocking buffer (1 mg/ml reduced BSA) for 1.5 h at RT. After 5 washes, the wells were incubated in 50 µl of primary antibody (anti-goat DNP 1:1000, A150-117A, Bethyl Laboratories) for 1 h at 37 °C, washed 5 times and incubated in 50 µl of secondary antibody (donkey anti-goat 1:2000, A15999, Invitrogen) for 2 h at 37 °C. After the final rounds of washing, the wells were incubated for 10-15 min at 37 °C in 50 µl with an o-phenylenediamine dihydrochloride (OPD, 34005, Thermo Scientific) solution (0.6 mg/ml OPD with 4 mmol H_2_O_2_, 50 mmol Na_2_HPO_4_ (71643, Sigma-Aldrich) and 2 mmol citric acid (5949-29-1, Fisher) in H_2_O). Color development was halted by addition of 30 µl of 2.5 M sulphuric acid (07208, Fluka), and the absorbance was measured at 485 nm with a Hidex Sense well plate reader. The results were calculated from the standard curve, normalized according to the protein content and to the mean of the WT group.

### Measuring protein carbonylation with immunoblotting

Protein carbonylation was additionally assessed with immunoblotting after incubation with DNP [79]. Single retinas were homogenized in 70 µl of 1XRIPA with 5 mM EDTA and 100 mM β-ME and centrifuged at 10 000 rpm for 10 min at 4 °C. Nucleic acids were removed, and an aliquot was taken for the protein determination with the Pierce BCA protein assay (23250, Thermo Scientific). Protein concentrations were equalized before DNP derivatization. For derivatization, 1 sample volume of 12 % (m/V) SDS (230425000, Thermo Scientific) was first added to the sample before adding 2 volumes of DNP (20 mM DNP in 10 % (V/V) trifluoroacetic acid (T/3258/04, Fisher Chemical)) or blank (10 % (V/V) trifluoroacetic acid) solution, and the sample was incubated in RT for 10 min, which was neutralized with 2 M TRIS (T2516, TCI) in 30 % (V/V) glycerol (G/0650/17, Fisher Scientific) solution. Derivatization was followed by electrophoresis at 150 V for 45 min with precast 4-15 % gels (4568085, Bio-Rad), after which the proteins were transferred to the PVDF membrane (Trans-Blot Turbo Midi 0.2 µm PVDF Transfer Packs, 1704157, Bio-Rad), blocked with 5 % (m/V) BSA in PBST and washed 3 times with PBST. Membranes were incubated overnight at 4 °C with the primary antibody (anti-goat DNP 1:10000, A150-117A, Bethyl Laboratories) and on the next day after washing 3 times with PBST secondary HRP antibody (donkey anti-goat 1:10000, A15999, Invitrogen) in RT for 2 h. Membranes were washed 3 times with PBST before rinsing finally with MilliQ water, and the chemiluminescence was induced as described earlier. The Azure Biosystems 600 was used for imaging. Results were analyzed with ImageJ and normalized to the total protein signal assessed with Coomassie staining (1610436, Bio-Rad) and to the mean of WT group.

### Measurement of antioxidant capacity

Antioxidant capacity was assessed with 2,2-diphenyl-1-picrylhydrazyl (DPPH) method [80]. Frozen retinas were homogenized with a hand-held device in ice-cold 1XPBS in a volume according to tissue mass (40 µl per 1 mg tissue), and centrifuged for 10 min at 4 °C at 10 000 G. Supernatant was collected, and 40 µl of sample was mixed in a light-protected tube with 350 µl of 1XPBS and 350 µl of 0.1 mM DPPH solution in methanol (A456-212, Fisher Scientific). After an 1 h incubation at RT in the dark the mixture was pipetted onto a 96-well plate (200 µl per well), which was immediately measured with Hidex Sense well plate reader at 520 nm. Results were calculated as fold-change to the WT group from the formula: 1) absorbance (Abs) reduction-% = (Abs_blank_ – Abs_sample_) / Abs_blank_ x 100; 2) nmol DPPH scavenged / mL supernatant = (Abs reduction-% / 100) x 50 (DPPH concentration 50 µmol/L) x 17.5 (sample dilution factor in tube). The 0.1 mM DPPH solution was prepared fresh on the day of the experiment by first dissolving 2 mg DPPH (D4313, Tokyo Chemical Industry Co.) to 500 µl of methanol (10 mM DPPH) and then diluting it further 100X with methanol to get the 0.1 mM working solution. Ascorbic acid (A0537, Tokyo Chemical Industry Co.) solutions in 1XPBS (10 – 7.5 – 5 – 3 – 1 – 0.5 mM; reaction volume 3.5 µl of standard solution with 346.5 µl 1XPBS and 350 µl DPPH) were used as positive control in experiments.

### Lipidomics

Fresh retinas were dissected, weighed, snap-frozen and stored in liquid nitrogen after direct cervical dislocation. Retina samples were thawed and homogenized on ice in Milli-Q water (1:20, W/V) using a handheld homogenizer for three cycles of 20 s, with 10-s intervals on ice. A total of 20 µl of retinal homogenates was transferred to new Eppendorf tubes while 20 µl of water was added to blank and double blank tubes. A total of 5 µl of lipid deuterated standard mixture (Splash Lipidomix Mass spec standard, 330707, Avanti) was added to each tube except for the double blank tubes. Lipids were extracted using methanol (A456-212, Fisher Scientific) and methyl tert-butyl ether (MTBE; G6529, Sigma-Aldrich) as previously described [81]. Briefly, 150 µl of ice-cold methanol was added to each tube which were then vortexed well, followed by addition of 500 µl of ice-cold MTBE; finally, they were incubated for one hour while mixing at 800 rpm at RT. A total of 125 µl of water was added to each tube and incubated for 10 min in RT during which time they were vortexed twice. The samples were then centrifuged at 1700 G for 10 minutes at 4 °C. The samples were then transferred into a dried-ice box for 15 minutes to freeze the lower aqueous phase while extracting the upper organic phase into new tubes. The extraction steps were repeated one more time using 645 µl of MTBE, 194 µl methanol, and 161 µl of water. The two organic phases (containing lipids) were combined and evaporated in speedvac for 1 hour at 30 °C. The samples were then stored at −80 °C until the day of analysis.

Lipid samples were resuspended in 100 µl of isopropanol (34863, Sigma-Aldrich) while mixing at RT for 15 min prior to filtration using Thermo Microcentrifugal units (F2517-9, Thermo Scientific). A total of 5 µl was analyzed by UHPLC (Vanquish Flex, Thermo Scientific) coupled to a high-resolution Orbitrap Q Exactive Classic mass spectrometer (Thermo Scientific) operating in positive and negative ion modes. The lipid samples were separated on Cortecs UPLC C18 column (2.1 × 100mm × 1.6 µm, Waters Co.) using mobile phase A: Acetonitrile (A955-212, Fisher Scientific): MilliQ water (50:50) supplemented with 10 mM ammonium formate (70221, Sigma-Aldrich) and 0.1% formic acid (70221, Sigma-Aldrich), and mobile phase B: Isopropanol: Acetonitrile: MilliQ water (88:10:2) supplemented with 2 mM ammonium formate and 0.1% formic acid. The flow rate was set at 0.3 ml/min, and lipids were separated over a 35-min gradient of 0-26 from 25-100 B followed by a four-minute washing period of 100% B prior to re-equilibration again for 5 minutes at 25% B. Data was acquired in the Full MS mode with a resolution of 140,000, AGC target 1e6, and a maximum IT 100 ms over a scan range of 120 to 1700 m/z, and data-dependent MS/MS mode with a resolution of 17,500 AGC target 5e5, maximum IT 50 ms, loop count 8, TopN 8, an isolation width of 1.2 m/z over a scan range of 200 to 2000 m/z.

The spectra RAW data was searched, aligned, and normalized using LipidSearch 5 (Thermo Scientific). Lipids were identified following the default Product Search for mammalian lipids module (precursor tolerance 5 ppm and product tolerance 10 ppm). Lipids were aligned across samples using 0.05 min as retention time tolerance. Lipid species from the phosphatidylcholine, phosphatidylethanolamine, phosphatidylserine, phosphatidylglycerol, phosphatidylinositol, phosphatidic acid, lysophosphatidylcholine, lysophosphatidylethanolamine, diglyceride, triglyceride, sphingomyelin and cholesteryl ester classes were normalized to their respective deuterated Splash lipid standards (Supplemental Table S4), while lipid species from ceramide, primary fatty acid amide, monoacylglycerol, sphinogosine base, coenzyme Q, wax ester, lysobisphosphatidic acid, and acylcarnitine groups were normalized to the deuterated Splash lipid standard PG(15:0_18:1)+NH4+D7. Samples with a 100 % identification rate in grade A and B were used for further downstream analysis and statistical lipid group comparisons. Only grade A identifications were used in the analysis of lipids with 20:4, 22:5 or 22:6 sidechains. For grade A identifications, all the class-specific ions and substituent-specific ions that define the structure were assigned, while for grade B identification, one of the class-specific ions and substituent-specific ions that designate the structure was assigned.

### RNA-seq analysis of open data

The bulk RNA sequencing data of healthy human retina was downloaded from the European Bioinformatics Institute (EBI) ArrayExpress (E-MTAB-4377) [41]. RNA-seq data analysis was performed following the general pipeline for quality control, alignment, and differential expression analysis. The quality of the raw RNA-seq reads was assessed using FastQC v0.11.9 [82] to identify potential issues with sequence quality, GC content, and adapter contamination. The alignment of the reads to the human genome was performed using STAR v2.7.11b [83] with the GRCh38 primary assembly genome file (GRCh38.primary_assembly.genome.fa) and the GENCODE v29 annotation file (gencode.v29.annotation.gtf). Feature counting was carried out using HTSeq v2.0 [84] to generate a read count matrix for each sample. For normalization and differential expression analysis, the generated count data was processed with R scripts, utilizing DESeq2 [85] for statistical analysis.

### Statistical analysis

Results were analyzed in GraphPad Prism (version 10.4, GraphPad Software) and are presented as mean ± SD. The normality of the data was determined with Shapiro-Wilk test, and the statistical significance was assessed subsequently with parametric (t-test, ordinary one-way analysis of variance (ANOVA) with Tukey’s, Brown-Forsythe ANOVA test with Dunnett’s T3, two-way ANOVA with Šídák’s, repeated measures two-way ANOVA or mixed-effects analysis with Šídák’s post-hoc) or with nonparametric statistical tests (Kruskal-Wallis with Dunn’s multiple comparisons). Statistical significance was set at p < 0.05, except in RNA-data where an adjusted p < 0.1 was used. Source data is presented in Supplemental Data 1 to 3.

## Data Availability

All study data are included in the article and/or supporting information. All code and analysis scripts are available in the public GitHub repository (https://github.com/UmairSeemab/RNAseqData).

## Supporting information

Supplemental Data 3 open human retina RNA-seq

Supplemental Data 2 Lipidomics data

## Acknowledgements

We thank Dr. Ewen MacDonald for proofreading and editing the manuscript. We thank Dr. Thierry Leveillard (Institut de la Vision, INSERM, France) for providing rd10 mice to establish our own colony. The use of Leica Slidescanner was made possible thanks to the UEF Cell and Tissue Imaging Unit, University of Eastern Finland, Biocenter Kuopio, and EuroBioimaging Finland, and the use of the QuantStudio 5 qPCR device was made possible thanks to Genome Center of Eastern Finland, University of Eastern Finland, Biocenter Kuopio, and EuroBioimaging Finland. We acknowledge the assistance of Merja Häkkinen, Seppo Auriola, and Marko Lehtonen for their guidance on LC-MS lipidomics and for their previous work on method optimization. This work was supported by grants from the Mary and Georg C. Ehrnrooth Foundation (Mary och Georg C. Ehrnrooths Stiftelse) and the Päivikki and Sakari Sohlberg Foundation (Päivikki ja Sakari Sohlbergin Säätiö). H.L. is additionally supported by grants from the Research Council of Finland (grant 346295), Emil Aaltonen Foundation (Emil Aaltosen Säätiö), Sigrid Jusélius Foundation (Sigrid Juséliuksen Säätiö), Finnish Eye and Tissue Bank Foundation (Silmä-ja kudospankkisäätiö), Retina Registered Association Finland (Retina ry), Sokeain Ystävät/De Blindas Vänner Registered Association, and the FEBS Excellence Award. K.V. is funded by the UEF Doctoral School.

## Author contributions

K.V. and H.L. designed the research; K.V., A.K., U.S., and A.B.M. performed the research; K.V. and A.B.M. contributed new reagents/analytic tools; K.V. and H.L. analyzed data; K.V and H.L. wrote the paper.

