## Supplementary Information for "Female sex is a risk factor for exacerbated lipid peroxidation and disease in murine retinitis pigmentosa"

**This file includes:**

Figures S1 to S14

Tables S1 to S4

Legends for Supplemental Data 1 to 3

SI References

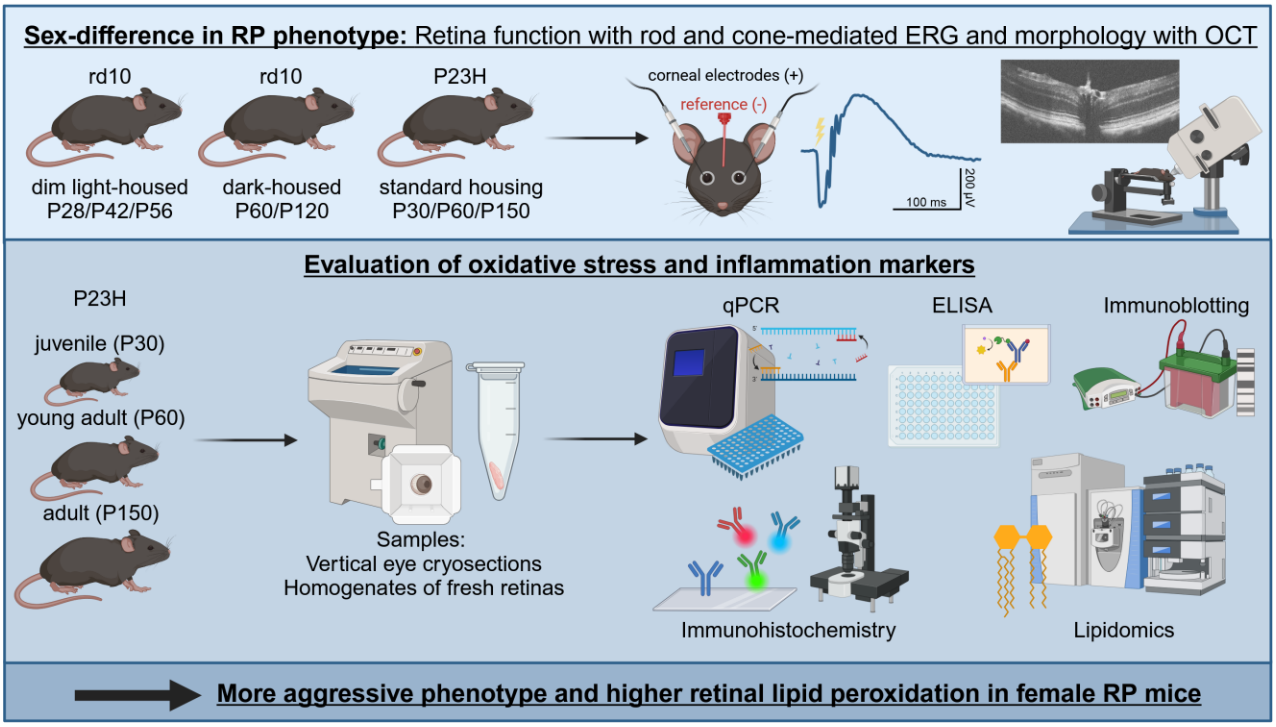


**Figure S1. Graphical abstract of the study.** Summary of used animal paradigms and methods.

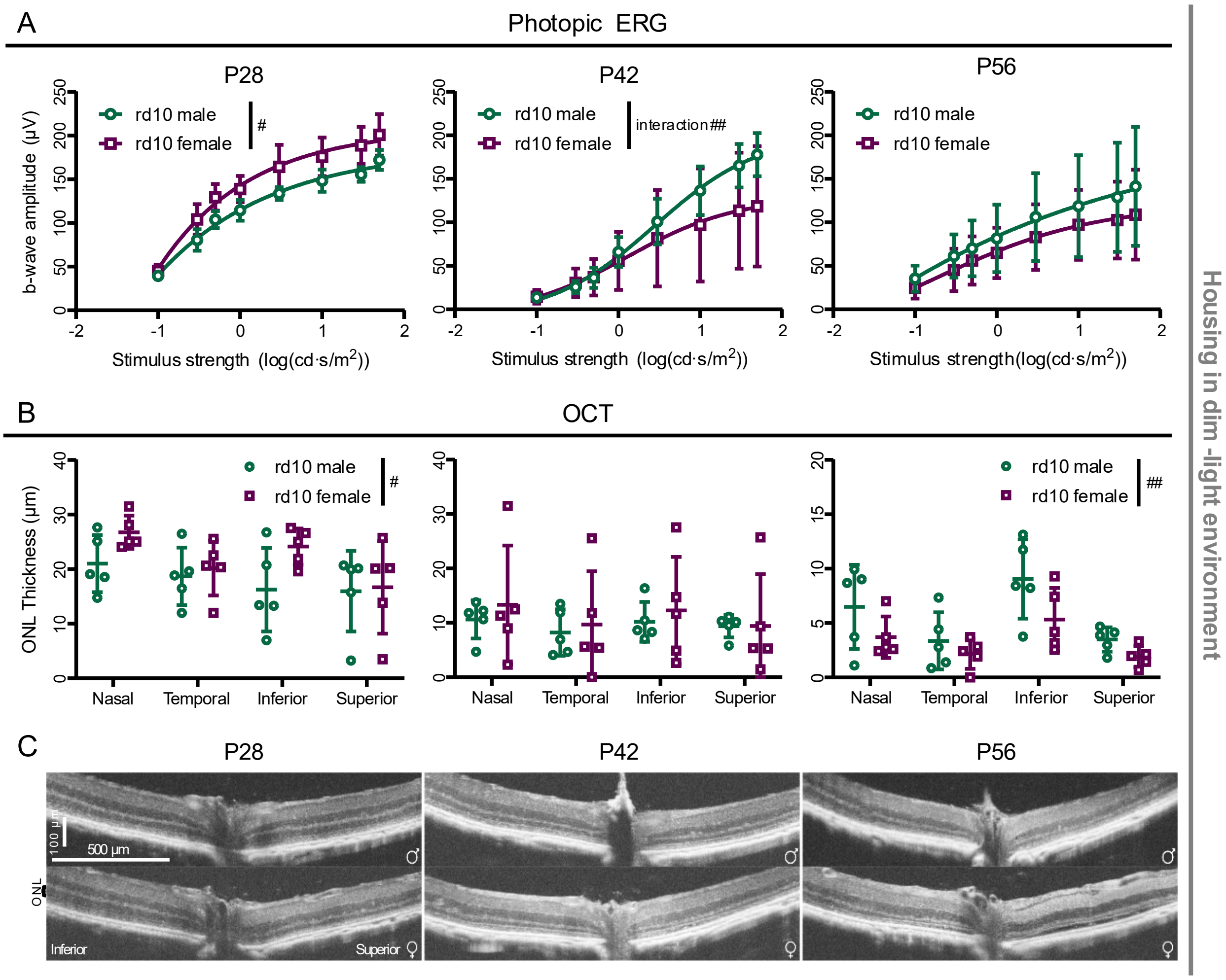


**Figure S2. Disease phenotype worsening in dim-light reared (< 1 lux) rd10 female mice coincides with sexual maturity.** A) Photopic electroretinogram (ERG) b-wave (blue stimulus), mean ± standard deviation (SD). P28, n = 5 males (m), n = 5 females (f); P42, n = 5 m, n = 4 f; P56, n = 5 m, n = 5 f. B) Outer nuclear layer (ONL) thicknesses at each timepoint, mean ± SD. C) Representative vertically oriented optical coherence tomography (OCT) images from each timepoint from male and female rd10 mice. The hashtags indicate significant between-subjects two-way ANOVA main effects (repeated measures (RM) ANOVA used in panel A and regular ANOVA in panel B): # = $p < 0.05$ and ## = $p < 0.01$.

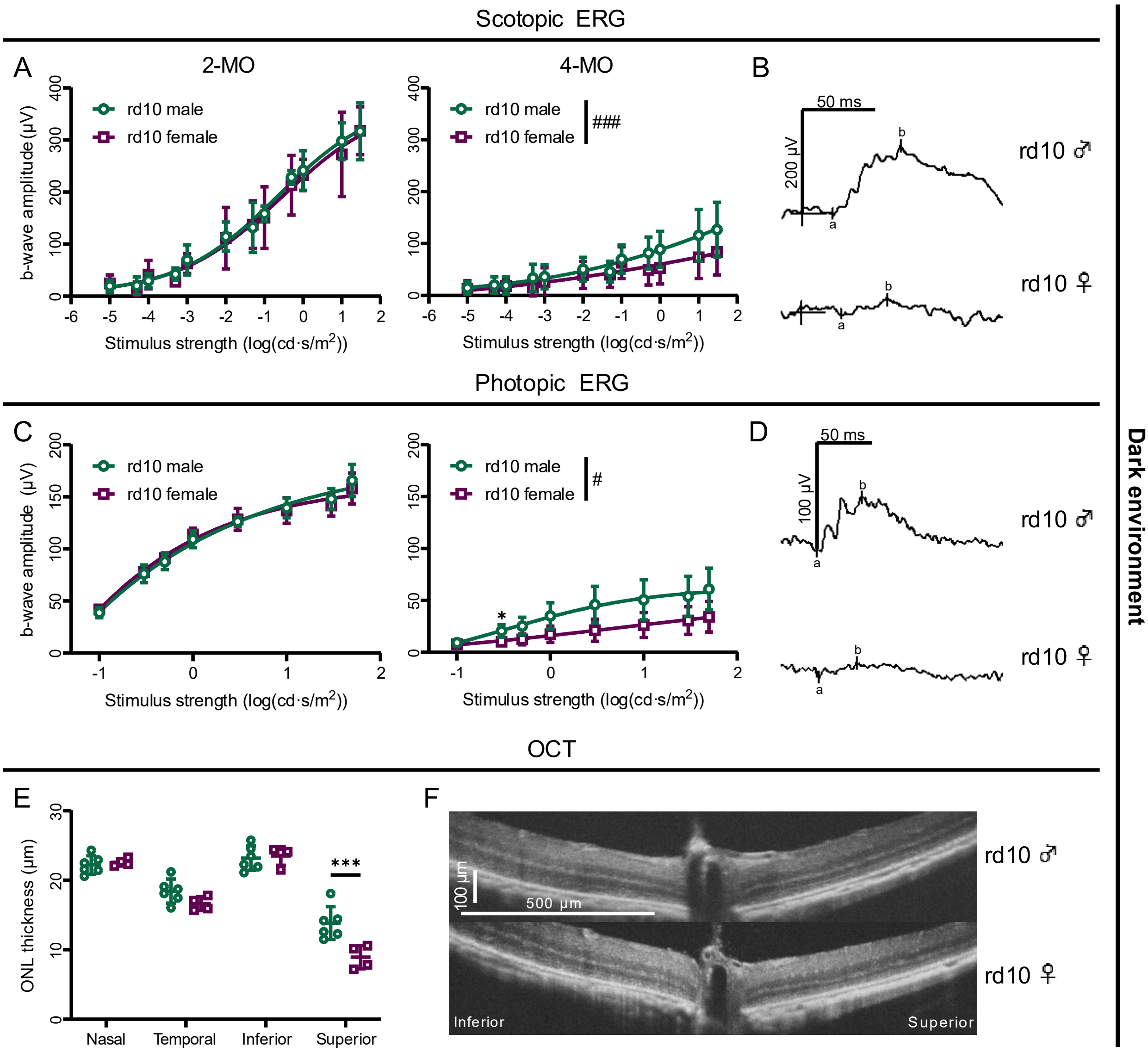


**Figure S3. Sex difference in adult rd10 mice is clear when constantly reared in full darkness.** This supplementary figure extends the data presented in main Figure 1. A) Scotopic ERG b-wave, mean ± SD. 2-month-old (MO), n = 6 m, n = 4 f; 4-MO, n = 15 m, n = 13 f. B) Representative scotopic ERG waveforms in 4-month-old mice (stimulus intensity: log 1 cd • s/m2). C) Photopic ERG b-wave (blue stimulus), mean ± SD. 2-MO, n = 6 m + 4 f; 4-MO n = 7 m + 6 f. D) Representative photopic ERG waveforms in 4-month-old mice (stimulus intensity: log 10 cd • s/m2). E) ONL thicknesses in 2-month-old mice, mean ± SD. F) Representative vertical OCT images from 2-month-old rd10 male and female mice. The hashtags and asterisks indicate significant between-subjects two-way ANOVA main effects (RM ANOVA used in panels A and C, and regular ANOVA in panel E) and Šídák's posthoc test results, respectively: #/* = $p < 0.05$, ##/** = $p < 0.01$ and ###/*** = $p < 0.001$.

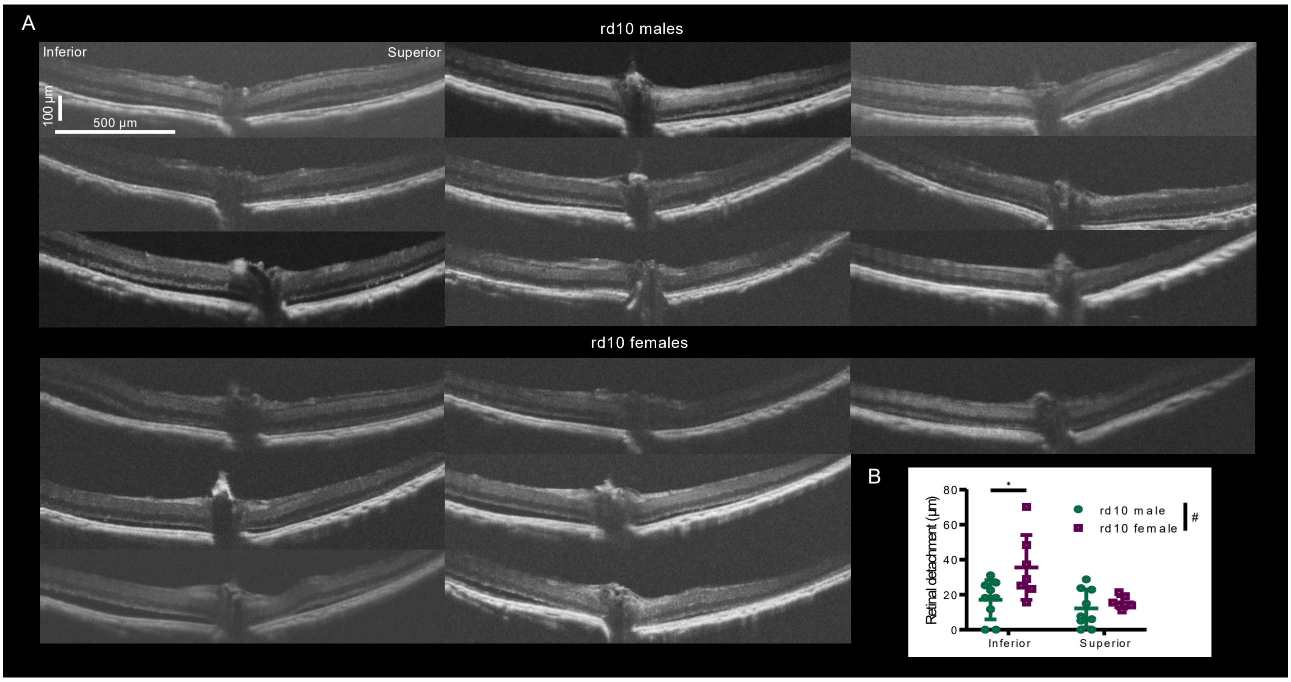


**Figure S4. Retinal detachment in the vertical retina is more severe in 4-month-old rd10 female mice.** This supplementary figure extends the data presented in main Figure 1. A) Vertically oriented example OCT images. B) Retinal detachment as measured from the highest point of detachment in inferior and superior orientations. The hashtags and asterisks indicate significant between-subjects ANOVA main effects and posthoc test results, respectively: #/* = $p < 0.05$, mean ± SD.

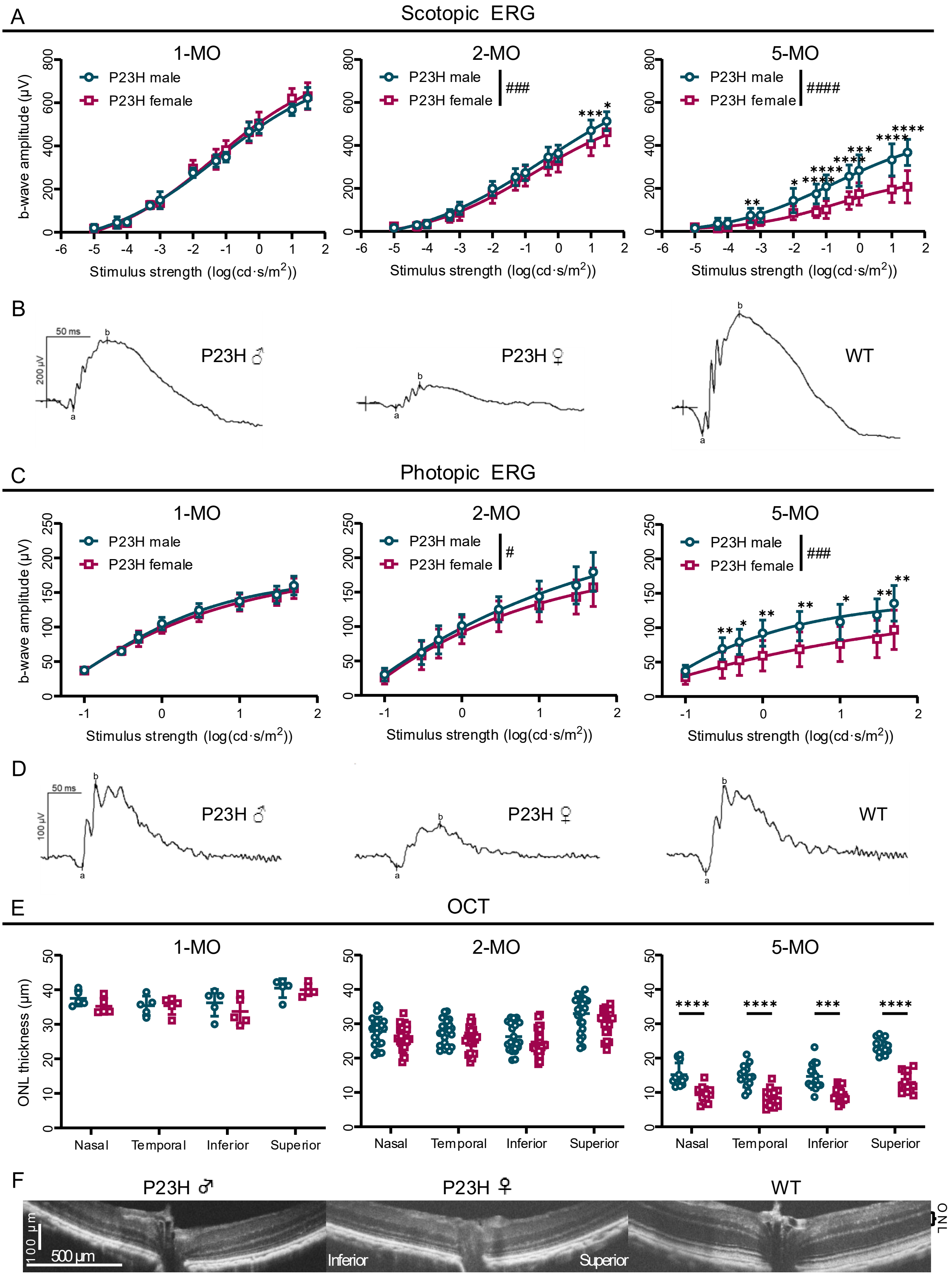


**Figure S5. Disease phenotype is accelerated in P23H female mice compared to males after puberty.** This supplementary figure extends the data presented in main Figure 1. A) Scotopic b-wave, mean ± SD. 1-MO, n = 5 m + 5 f; 2-MO, n = 24 m, n = 26 f; 5-MO, n = 16 m, n = 14 f. B) Representative scotopic ERG waveforms in 5-month-old mice (stimulus intensity: log 0.05 cd • s/m2). C) Photopic ERG b-wave (blue stimulus), mean ± SD. 1-MO, n = 5 m, n = 5 f; 2-MO, n = 24 m, n =

26 f; 5-MO, n = 15 m, n = 14 f. D) Representative photopic ERG waveforms in 5-month-old mice (stimulus intensity: log 10 cd • s/m2). E) ONL thicknesses, mean ± SD. F) Representative vertical OCT images from 5-month-old male and female P23H, and WT mice. The hashtags and asterisks indicate significant between-subjects two-way ANOVA main effects (RM ANOVA used in panels A and C, and regular ANOVA in panel E) and Šídák's posthoc test results, respectively: #/* = $p < 0.05$, ##/** = $p < 0.01$, ###/*** = $p < 0.001$ and ####/**** = $p < 0.0001$.

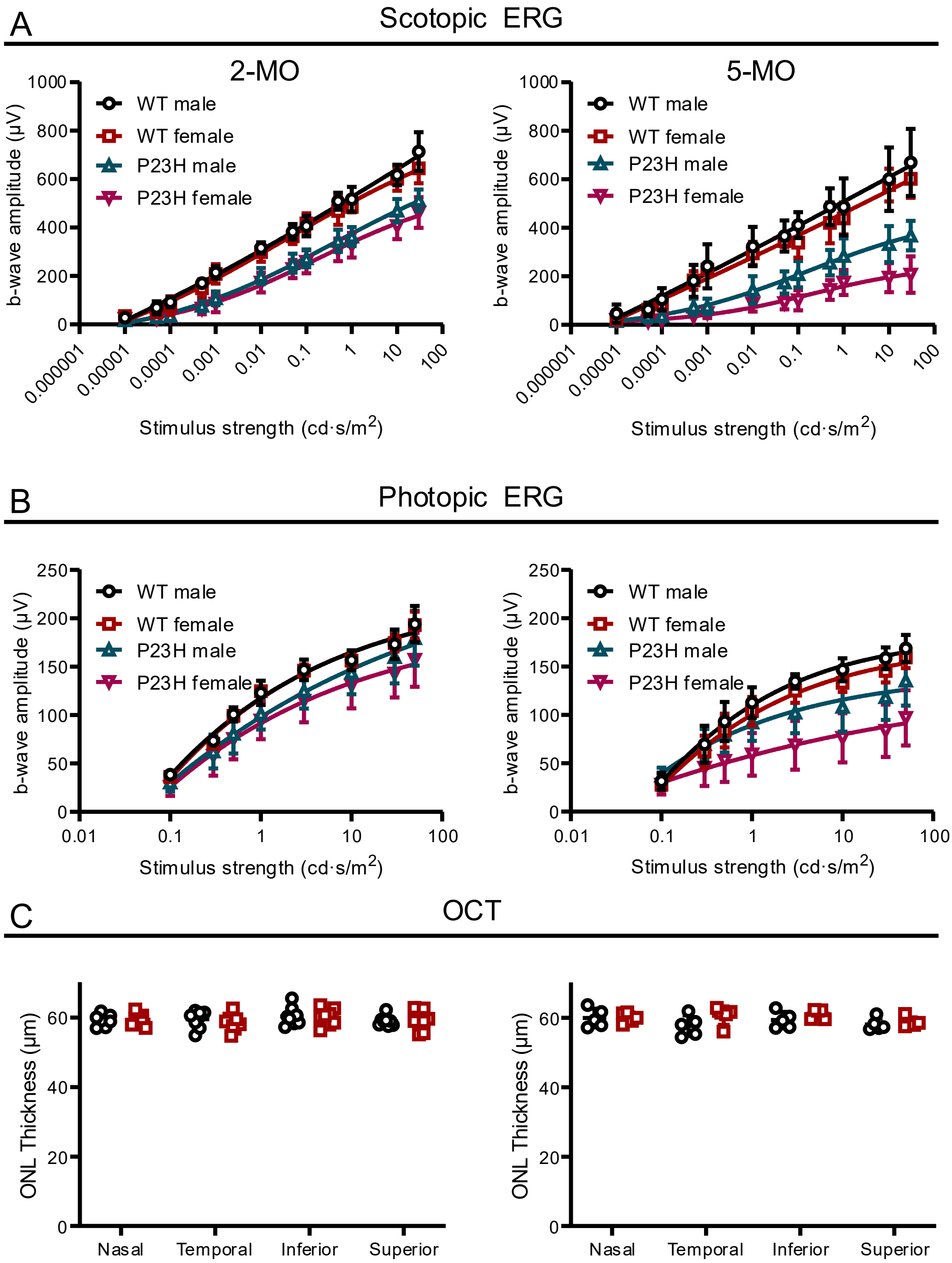


**Figure S6. WT mice display no sex differences in retinal function or thickness.** A-B) ERG amplitudes and C) ONL thickness in WT female versus male mice. WT 2-MO, n = 8 m, n = 8 f, in scotopic ERG and OCT, and n = 3 m, n = 3 f, in photopic ERG. WT 5-MO, n = 5 m, n = 5 f, in all experiments. For comparison, P23H ERG data here is reproduced from Supplemental figure S5. Data is presented as mean ± SD.

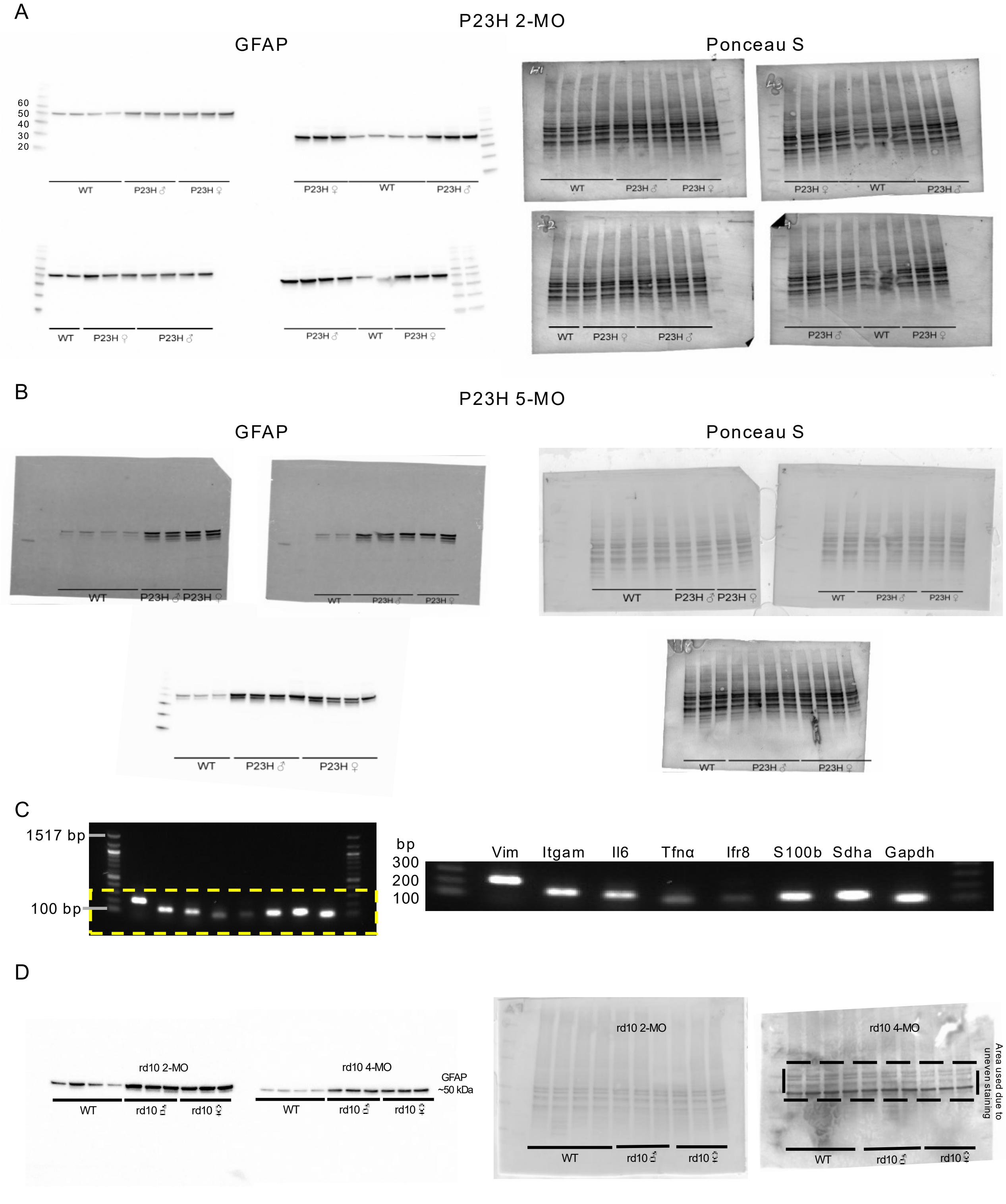


**Figure S7. Full immunoblots (A,B), and endpoint PCR with primers used in qPCR.** This figure relates to main text Figures 2 (A-C) and 4 (D). All analysed glial fibrillary acidic protein (GFAP) and Ponceau S stained immunoblots with A) 2-month-old and B) 5-month-old P23H and WT retina samples. C) Electrophoresis with example sample after amplification with vimentin (*Vim*), integrin subunit alpha M (*Itgam*), interleukin 6 (*Il6*), tumor necrosis factor alpha (*Tnfa*), interferon regulatory factor 8 (*Irf8*), S100 calcium-binding protein B (*S100b*), succinate dehydrogenase complex flavoprotein subunit A (*Sdha*) and glyceraldehyde 3-phosphate dehydrogenase (*Gapdh*) primers. D) All analysed GFAP and Ponceau S stained immunoblots with rd10 and WT retina samples.

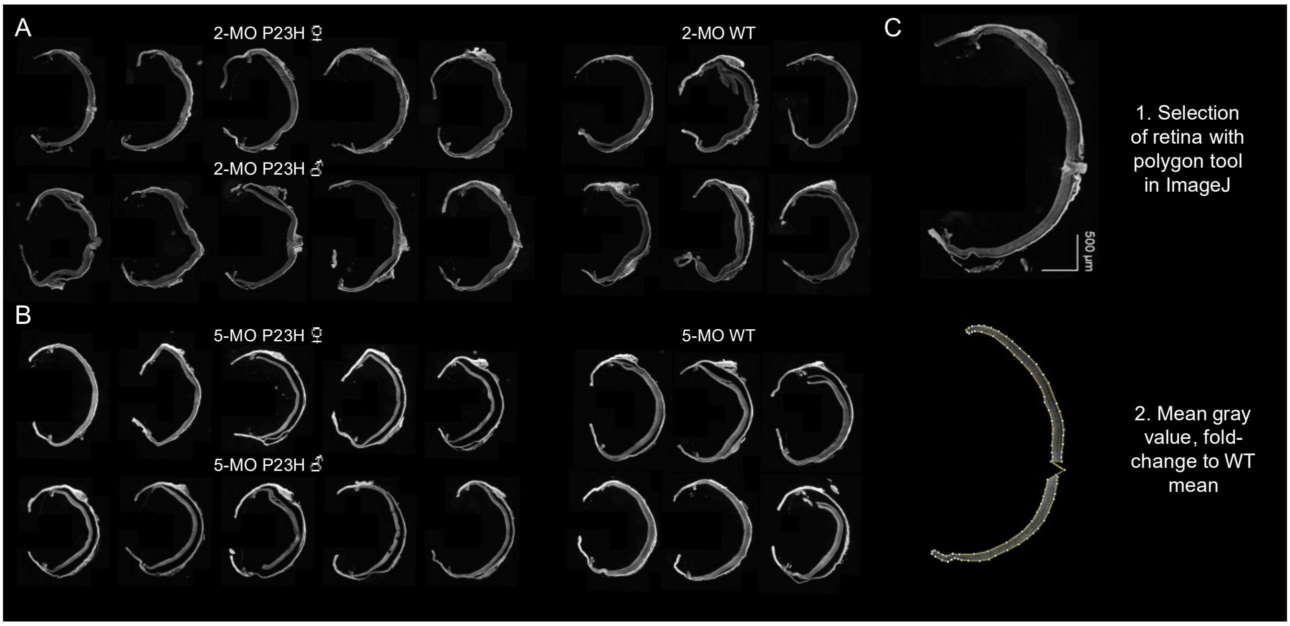


**Figure S8. 4-HNE signal from all samples used for analysis in main text Figure 3B.** 4-HNE-stained cryosections from vertically oriented eye cups from A) 2- and B) 5-month-old P23H and WT mice. C) Example of retinal area selection for quantification.

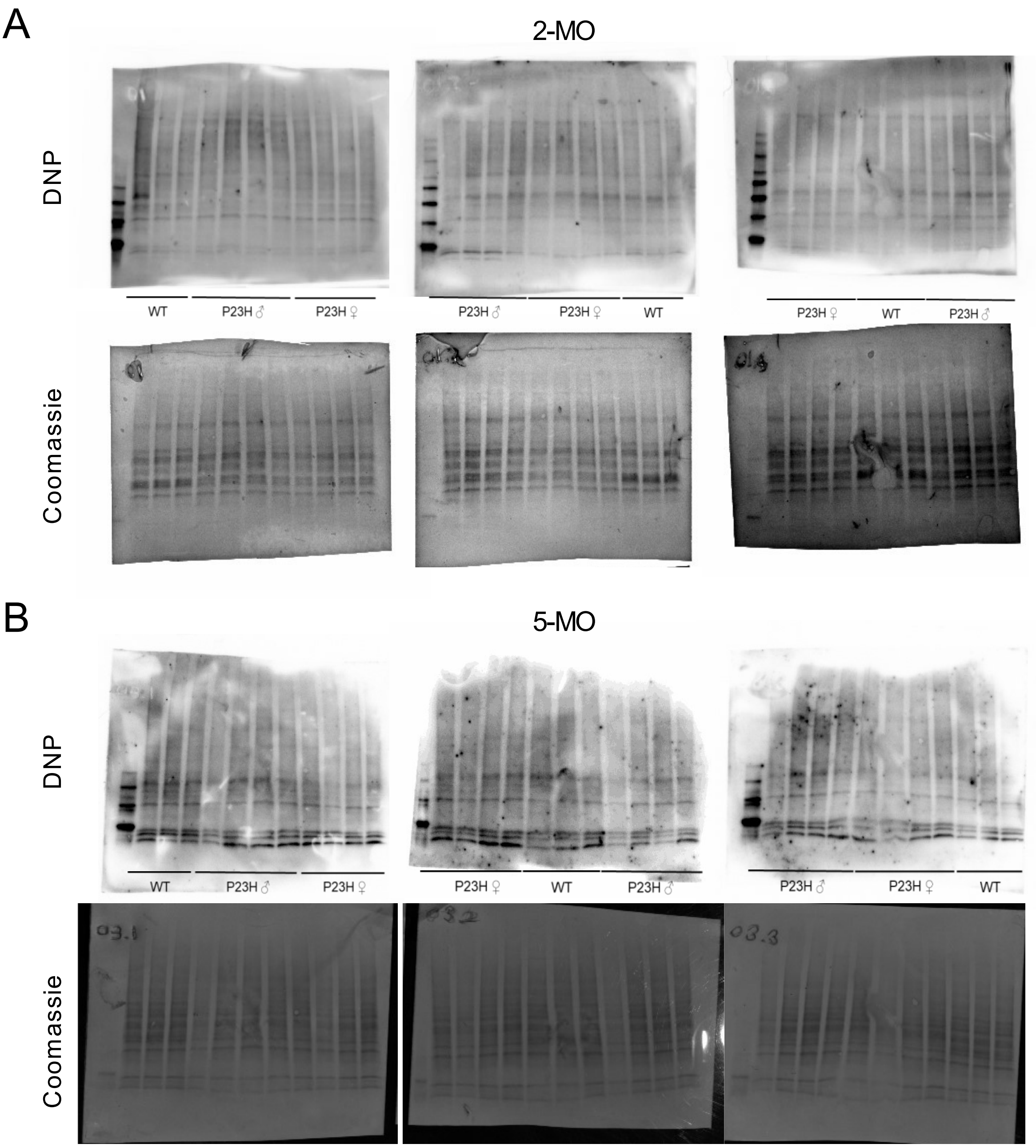


C

1. Lane selection with lane tool in ImageJ

2. Plotting the lanes and measurement of area under the peaks with high background reducement

3. Sum of AUCs, normalization to Coomassie AUC and fold-change to WT mean

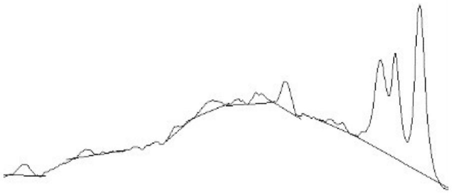

**Figure S9. Measurement of protein carbonylation by immunoblotting.** Relates to main text Figure 3E. All analysed 2,4-dinitrophenylhydrazine (DNP) and Coomassie stained blots with A) 2- and B) 5-month-old P23H and WT retina samples. C) Workflow for image analysis.

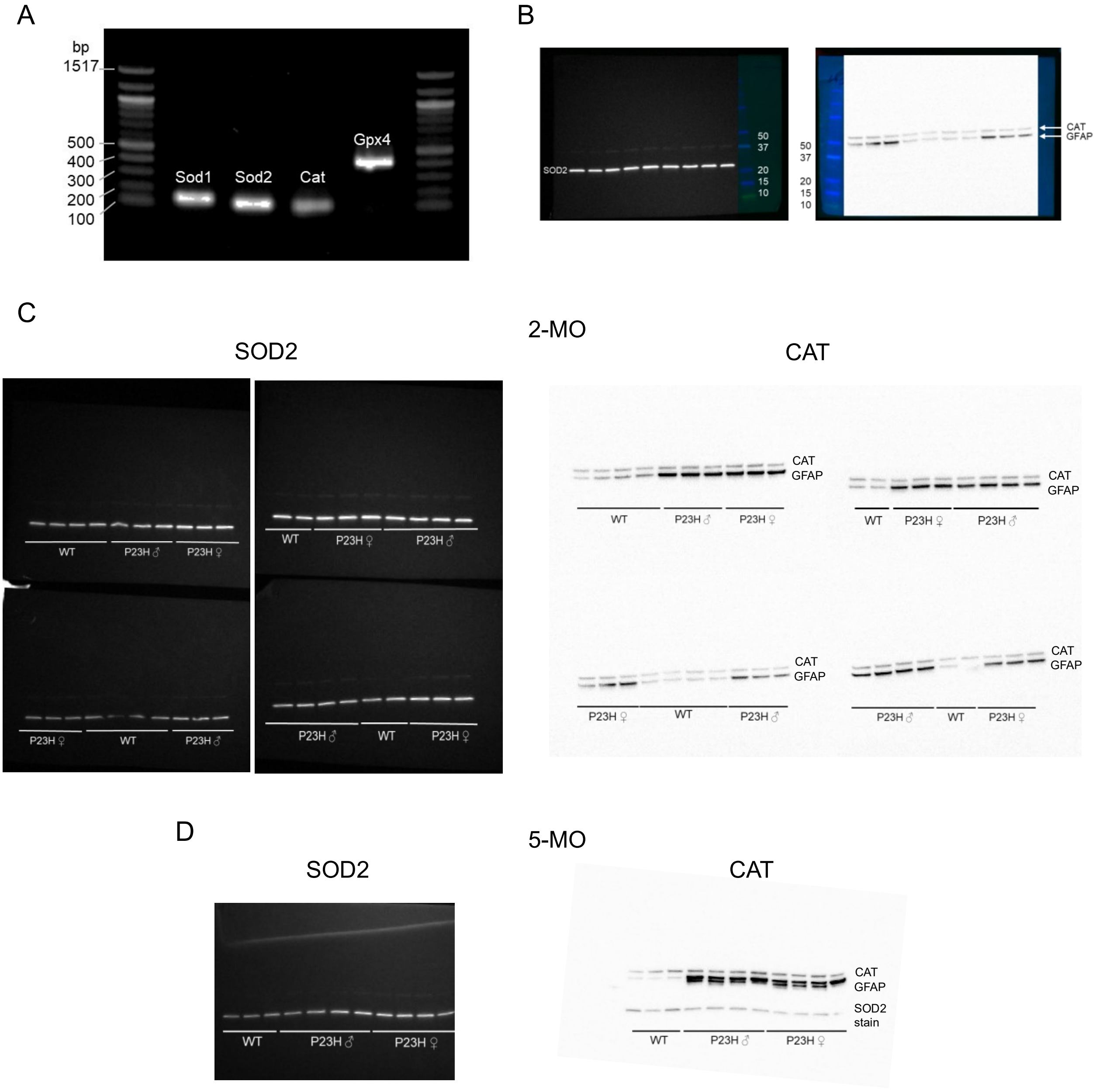


**Figure S10. Assessment of antioxidative system in male and female P23H retinas.** A) Electrophoresis with example samples after amplification with superoxide dismutase 1 (*Sod1)*, superoxide dismutase 2 (*Sod2*), catalase (*Cat*) and glutathione peroxidase 4 (*Gpx4*) primers. Relates to main text Figures 3G-J. B) Merge images of membranes with SOD2 and CAT (and glial fibrillary acidic protein (GFAP) remnant) signals with molecular weight markers. Full images of immunoblots stained with SOD2 and CAT with retina samples from C) 2- and D) 5-month-old P23H and WT mice. Panels B-D relate to main text Figures 3K-M.

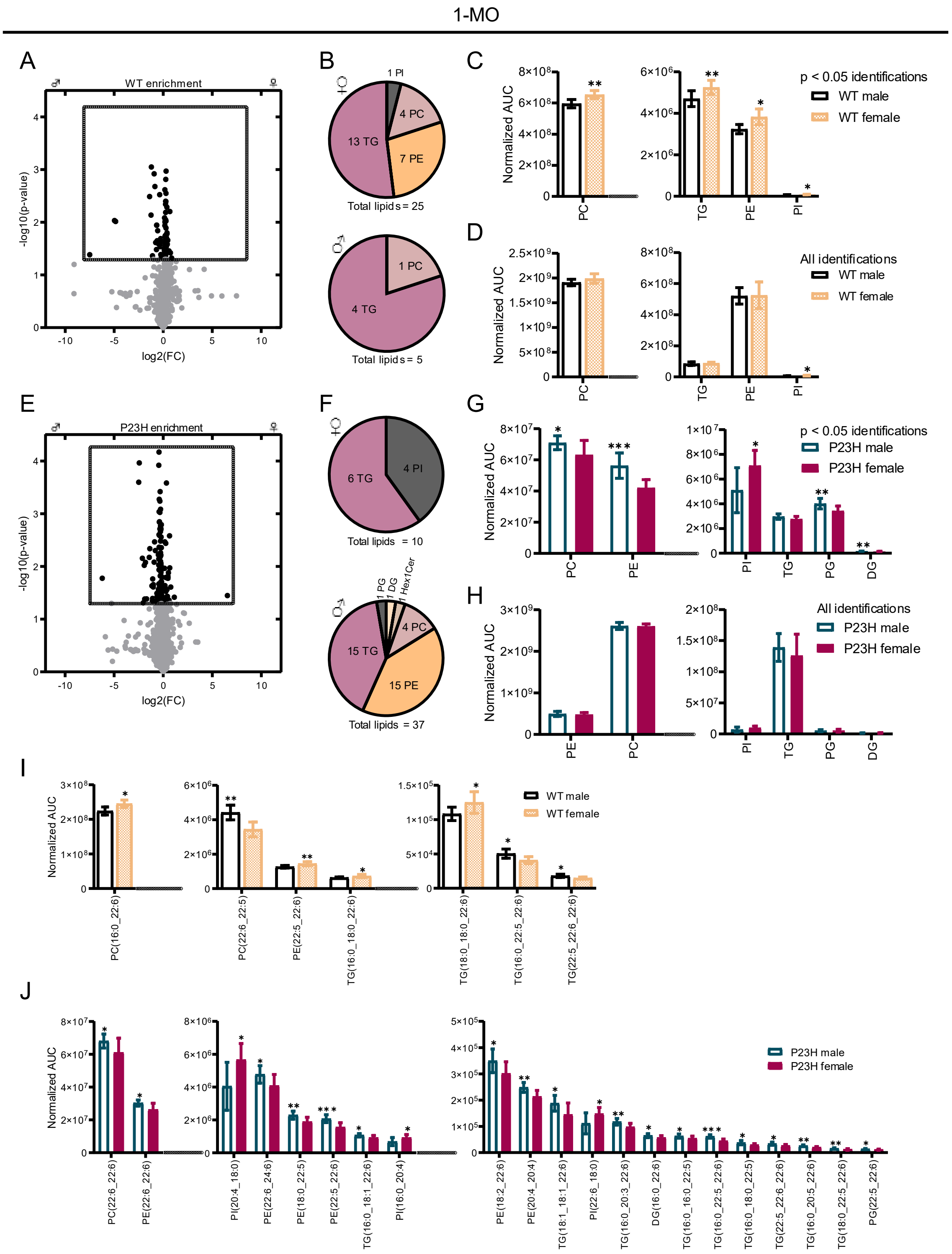


**Figure S11. Comparison of retinal lipids between 1-month-old WT male and WT female, and P23H male and P23H female mice.** A) Volcano plot of all identified lipids in WT mice. Differentially expressed lipids (DELs, $p < 0.05$) between WT males ($n = 7$) and females ($n = 5$) are shown in black. B) Number of enriched DELs per lipid class in WT females and males. C) Lipid group expression comparison between WT male and female mice based on DELs only. D) Lipid group expression comparison between WT male and female mice based on all lipid identifications. E) Volcano plot of all identified lipids in P23H mice ($n = 9$ males; $n = 8$ females). F) Number of enriched DELs per lipid class in P23H females and males. G) Lipid group expression comparison between P23H male and female mice based on DELs only. H) Lipid group expression comparison between P23H male and female mice based on all lipid identifications. I) DELs with 20:4/22:5/22:6 carbon/double-bond sidechains between WT male and female mice, and J) between P23H male and female mice. In all graphs, the asterisks indicate significant t-test results, * = $p < 0.05$, ** = $p < 0.01$ and *** = $p < 0.001$. The bar graph data is presented as mean ± SD. DG = diglyceride, PC = phosphatidylcholine, PE = phosphatidylethanolamine, PI = phosphatidylinositol, PG = phosphatidylglycerol, TG = triglyceride.

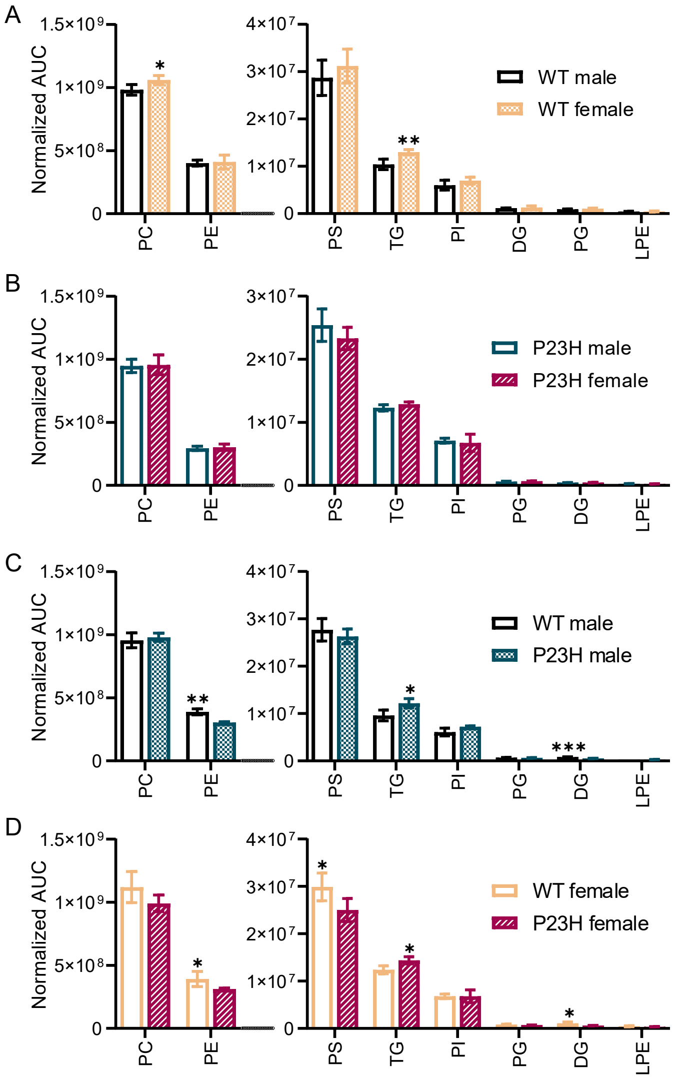


**Figure S12. Lipid group expression comparisons with all grade A lipid identification with LC-PUFA (20:4, 22:5 or 22:6) sidechains, 2-month-old WT and P23H retinas.** Comparisons are between A) WT male (n = 4) and WT (n = 4) female, B) P23H male (n = 3) and P23H female (n = 4), C) WT male and P23H male, and D) WT female and P23H female retinas, mean ±SD. In all graphs, the asterisks indicate significant t-test results, * = $p < 0.05$, ** = $p < 0.01$ and *** = $p < 0.001$. DG = diglyceride, LPE = lysophosphatidylethanolamine, PC = phosphatidylcholine, PE = phosphatidylethanolamine, PG = phosphatidylglycerol, PI = phosphatidylinositol, PS = phosphatidylserine, TG = triglyceride.

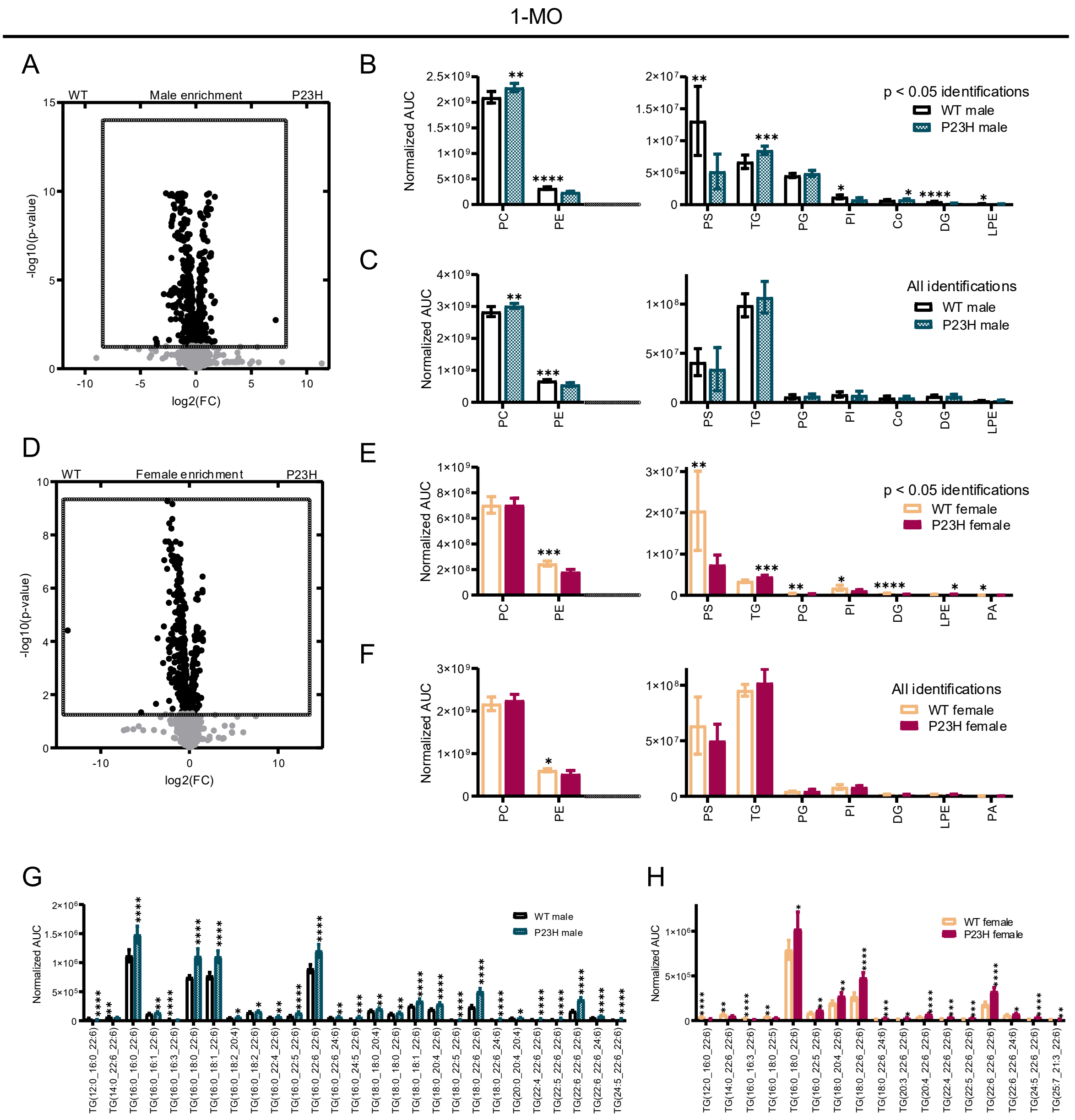


**Figure S13. Analysis of differentially expressed lipids between genotypes for both sexes at the age of 1 month.** A) Volcano plot of all identified lipids in WT and P23H males. Differentially expressed lipids (DELs) between WT males (n = 7) and P23H males (n = 9) are shown in black. B) Lipid group expression comparison between WT males and P23H males based on DELs only. C) Lipid group expression comparison between WT males and P23H males based on all lipid identifications. D) Volcano plot of all identified lipids in WT and P23H females. DELs between WT females (n = 5) and P23H females (n = 8) are shown in black. E) Lipid group expression comparison between WT females and P23H females based on DELs only. F) Lipid group expression comparison between WT females and P23H females based on all lipid identifications. G) Differentially expressed TGs with 20:4/22:5/22:6 carbon/double-bond sidechains between WT males and P23H males, and H) between WT females and P23H females. In all graphs, the asterisks indicate significant t-test results, * = $p < 0.05$, ** = $p < 0.01$, *** = $p < 0.001$ and **** = $p < 0.0001$. The bar graph data is presented as mean ±SD. Co = coenzyme Q, DG = diglyceride, LPE = lysophosphatidylethanolamine, PA = phosphatidic acid, PC = phosphatidylcholine, PE = phosphatidylethanolamine, PG = phosphatidylglycerol, PI = phosphatidylinositol, PS = phosphatidylserine, TG = triglyceride.

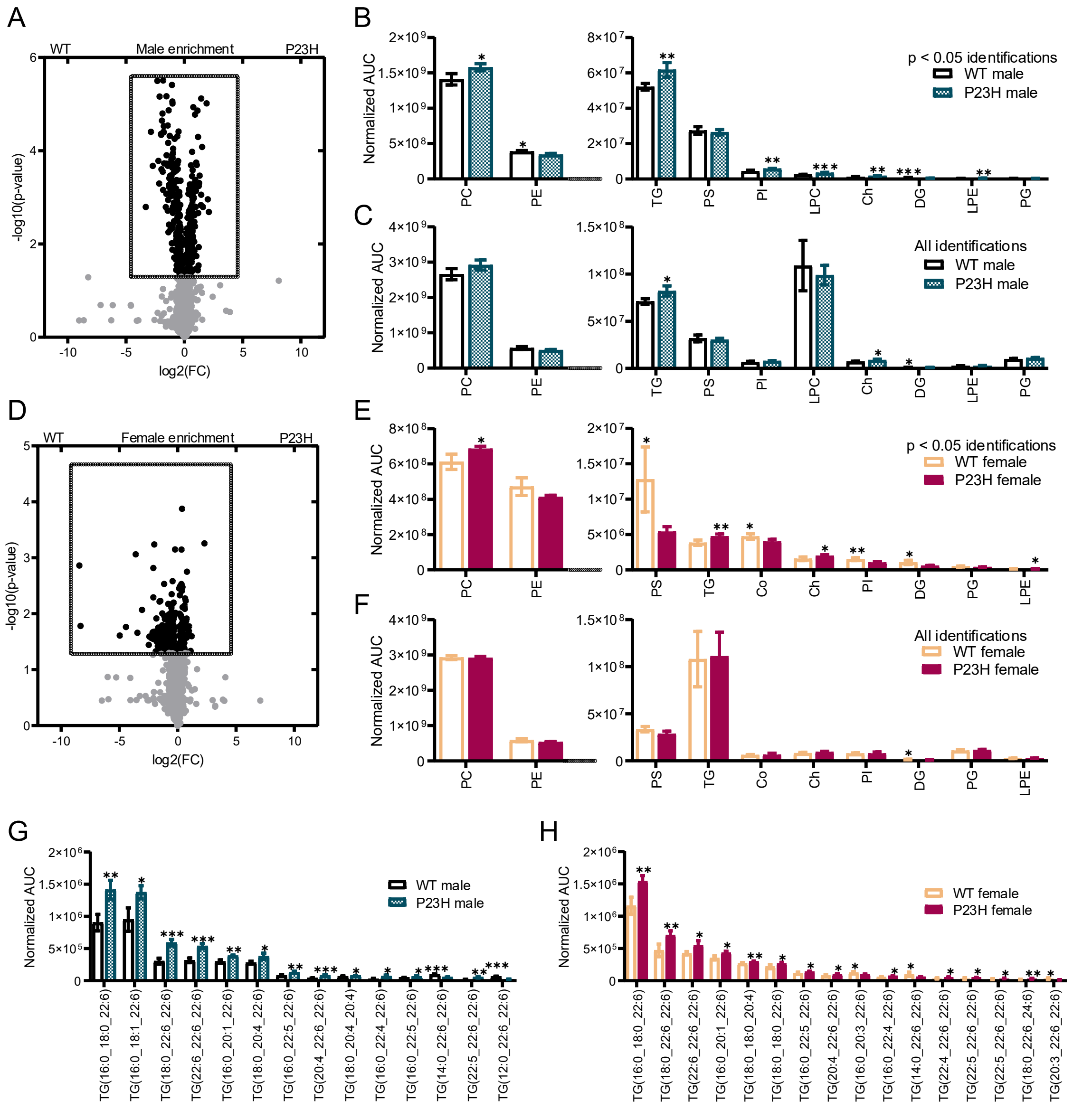


**Figure S14. Analysis of differentially expressed lipids between genotypes for both sexes at the age of 2 months.** A) Volcano plot of all identified lipids in WT and P23H males. Differentially expressed lipids (DELs) between WT males (n = 4) and P23H males (n = 3) are shown in black. B) Lipid group expression comparison between WT males and P23H males based on DELs only. C) Lipid group expression comparison between WT males and P23H males based on all lipid identifications. D) Volcano plot of all identified lipids in WT and P23H females. DELs between WT females (n = 4) and P23H females (n = 4) are shown in black. E) Lipid group expression comparison between WT females and P23H females based on DELs only. F) Lipid group expression comparison between WT females and P23H females based on all lipid identifications. G) Differentially expressed TGs with 20:4/22:5/22:6 carbon/double-bond sidechains between WT males and P23H males, and H) between WT females and P23H females. In all graphs, the asterisks indicate significant t-test results, * = $p < 0.05$, ** = $p < 0.01$ and *** = $p < 0.001$. The bar graph data is presented as mean ±SD. Co = coenzyme Q, Ch = Cholestrol, DG = diglyceride, LPC = lysophosphatidylcholine, LPE = lysophosphatidylethanolamine, PC = phosphatidylcholine, PE = phosphatidylethanolamine, PG = phosphatidylglycerol, PI = phosphatidylinositol, PS = phosphatidylserine, TG = triglyceride.

**Table S1. Protocol for scotopic electroretinography stimulation.** ISI, inter-stimulus interval time.

| Step | Stimulus (cd•s/m$^2$), green | ISI (s) | Repetitions |
|---|---|---|---|
| 1 | 0.00001 | 1 | 25 |
| 2 | 0.00005 | 2 | 20 |
| 3 | 0.0001 | 2 | 20 |
| 4 | 0.0005 | 3 | 15 |
| 5 | 0.001 | 5 | 12 |
| 6 | 0.01 | 10 | 8 |
| 7 | 0.05 | 10 | 8 |
| 8 | 0.1 | 15 | 4 |
| 9 | 0.5 | 20 | 4 |
| 10 | 1 | 20 | 3 |
| 11 | 10 | 45 | 2 |
| 12 | 30 | 60 | 2 |

**Table S2. Protocol for photopic electroretinography stimulation.**

| Step | Stimulus (cd•s/m$^2$), blue | ISI (s) | Repetitions |
|---|---|---|---|
| 1 | 0.1 | 0.15 | 40 |
| 2 | 0.3 | 0.15 | 30 |
| 3 | 0.5 | 0.15 | 25 |
| 4 | 1 | 0.5 | 15 |
| 5 | 3 | 0.5 | 15 |
| 6 | 10 | 0.75 | 10 |
| 7 | 30 | 0.75 | 10 |
| 8 | 50 | 1 | 10 |

**Table S3. Primers for qPCR.**

| Gene | Marker | Forward (Fw) – Reverse (Rv) |
|---|---|---|
| Vimentin (Vim) | Müller glia activation (1). | Fw: TTCTCTGGCACGTCTTGACC<br>Rv: CTCCTGGAGGTTCTTGGCAG |
| Integrin subunit alpha M (Itgam/Cd11b) | Microglia marker activation (2). | Fw: CTTTGGGAACCTCCGACCAG<br>Rv: CACCAAAGTGTCCAAGCCCA |
| Interleukin 6 (Il6) | Proinflammatory cytokine (3). | Fw: ATCCAGTTGCCTTCTTGGGACTGA<br>Rv: TAAGCCTCCGACTTGTGAAGTGGT |
| Tumour necrosis factor alpha (Tnfa) | Proinflammatory cytokine (4). | Fw: TCTCATGCACCACCATCAAGGACT<br>Rv: ACCACTCTCCCTTTGCAGAACTCA |
| Interferon regulatory factor 8 (Irf8) | Reactive microglia (5). | Fw: TTCAAGGCAGGTGGTGGT<br>Rv: GGATATGCCGCCTATGACAC |
| S100 calcium-binding protein B (S100b) | Gliosis (6). | Fw: CTGGAGAAGGCCATGGTTGC<br>Rv: CTCCAGGAAGTGAGAGAGCT |
| Superoxide dismutase 1 (Sod1) | Oxidative stress, conversion of superoxide radicals into hydrogen peroxide (7). | Fw: AACCAGTTGTGTTGTCAGGAC<br>Rv: CCACCATGTTTCTTAGAGTGAGG |
| Superoxide dismutase 2 (Sod2) | Oxidative stress, conversion of superoxide radicals into hydrogen peroxide (7). | Fw: CAGACCTGCCTTACGACTATGG<br>Rv: CTCGGTGGCGTTGAGATTGTT |
| Catalase (Cat) | Breakdown of hydrogen peroxide to water and oxygen (7). | Fw: GGAGGCGGGAACCCAATAG<br>Rv: GTGTGCCATCTCGTCAGTGAA |
| Glutathione peroxidase 4 (Gpx4) | Breakdown of hydrogen peroxide to water and oxygen, lipid peroxidation (7). | Fw: GATGGAGCCCATTCCTGAACC<br>Rv: CCCTGTACTTATCCAGGCAGA |
| Succinate Dehydrogenase Complex Flavoprotein Subunit A (Sdha) | Housekeeping gene 1 | Fw: GCTCCTGCCTCTGTGGTTGA<br>Rv: AGCAACACCGATGAGCCTG |
| Glyceraldehyde-3-Phosphate Dehydrogenase (Gapdh) | Housekeeping gene 2 | Fw: TGACGTGCCGCCTGGAGAAA<br>Rv: AGTGTAGCCCAAGATGCCCTTCAG |

**Table S4. Internal splash standards for phosphatidylcholine (PC), phosphatidylethanolamine (PE), phosphatidylserine (PS), phosphatidylglycerol (PG), phosphatidylinositol (PI), phosphatidic acid (PA), lysophosphatidylcholine (LPC), lysophosphatidylethanolamine (LPA), diglyceride (DG), triglyceride (TG), sphingomyelin (SM) and cholestryl ester (ChE) lipid groups.**

| Lipid group | Splash standard |
|---|---|
| PC | PC(33:1)+H+D7:(s) |
| PE | PE(15:0_18:1)+H+D7:(s) |
| PS | PS(33:1)+H+D7:(s) |
| PG | PG(15:0_18:1)+NH4+D7:(s) |
| PI | PI(18:1_15:0)+NH4+D7:(s) |
| PA | PA(15:0_18:1)+NH4+D7:(s) |
| LPC | LPC(18:1)+H+D7:(s) |
| LPE | LPE(18:1)+H+D7:(s) |
| DG | DG(33:1)+H+D7:(s) |
| TG | TG(15:0_15:0_18:1)+NH4+D7:(s) |
| SM | PG(15:0_18:1)+NH4+D7:(s) |
| ChE | Ch-D7+H-H2O |

**Supplemental Data 1 (separate file).** Raw data and animal information file.

**Supplemental Data 2 (separate file).** Lipidomics data file.

**Supplemental Data 3 (separate file).** Re-analysis of publicly available human bulk retina RNA-seq dataset.
